# *Platr4* is an ESC-specific lncRNA that exhibits its function downstream on meso/endoderm lineage commitment

**DOI:** 10.1101/2021.12.20.473435

**Authors:** Rasmani Hazra, Lily Brine, Libia Garcia, Brian Benz, Napon Chirathivat, Michael M. Shen, John Erby Wilkinson, Scott K. Lyons, David L. Spector

## Abstract

The mammalian genome encodes thousands of long non-coding RNAs (lncRNAs) that are developmentally regulated and differentially expressed across tissues, suggesting possible roles in cellular differentiation. Despite this expression pattern, little is known about how lncRNAs influence lineage commitment at the molecular level. Here, we reveal that perturbation of an embryonic stem cell (ESC)-specific lncRNA, *Pluripotency associated transcript 4* (*Platr4*), in ESCs directly influences the downstream meso/endoderm differentiation program without affecting pluripotency. We further show that *Platr4* interacts with the TEA domain transcription factor 4 (Tead4) to regulate the expression of a downstream target gene crucial in the cardiac lineage program known as connective tissue growth factor (Ctgf). Importantly, *Platr4* knockout mice exhibit myocardial atrophy, valve mucinous degenration associated with reduced cardiac output and sudden heart failure. Together, our findings provide evidence that *Platr4* expression in undifferentiated ESCs is critical for downstream lineage differentiation, highlighting its importance in disease modeling and regenerative medicine.

## Introduction

Recent advances in sequencing technologies have revealed that a large part of the human and mouse genomes encode for long ncRNAs (lncRNAs) which are greater than 200 nucleotides in length (Rinn and Chang, 2012; Wang and Chang, 2011). A significant number of lncRNAs are critical for ESC pluripotency and/or differentiation (Derrien et al., 2012; Guttman et al., 2011; Mercer et al., 2010), yet their regulatory role during these critical processes is not fully understood. Studies in ESCs have shown that complex biological networks, consisting of transcription factors, signaling pathways, and non-coding RNAs (ncRNAs), collaborate in a synergistic manner to regulate stemness and cell fate (Chen et al., 2020a). Many lncRNAs are expressed in a cell/tissue-specific and/or developmentally-specific manner, suggesting a possible role in lineage commitment and/or cellular differentiation.

Multiple studies have shown that lncRNAs are involved in lineage commitment and cell fate specification (Smith et al., 2019). A neural-specific lncRNA, *Pnky* regulates mouse and human neurogenesis by interacting with polypyrimidine tract-binding protein 1 (Ramos et al., 2015). Further, Eomes expressing mesendoderm progenitor lncRNA *Meteor* interacts with Eomes and epigenetically regulates mesendoderm specification and cardiac differentiation (Alexanian et al., 2017; Guo et al., 2018). In addition, *Braveheart* (*Bvht*), a heart-specific lncRNA, functions upstream of mesoderm posterior 1 (Mesp1) and regulates a cardiovascular gene network by interacting with SUZ12, a core component of the PRC2 complex (Klattenhoff et al., 2013). Despite these tissue/developmental-specific roles of lncRNAs, their function in ESCs during lineage specification and differentiation has yet to be investigated. Interestingly, cardiac-specific lncRNAs have been shown to be involved in cardiac function and diseases (Hobuss et al., 2019).

A cysteine-rich 38kD protein, connective tissue growth factor (CTGF), a member of the CCN family of matricellular protein, has been associated with maintenance of normal cardiac function and various cardiac diseases (Gerritsen et al., 2016; Leeuwis et al., 2010; Rickard et al., 2009), It has also been demonstrated that CTGF is critical for inducing the differentiation of many cell types; however, a role in cardiomyocyte (CM) specification has not yet been established (Croci et al., 2004; Luo et al., 2004; Morrison et al., 2010). CTGF is a direct downstream target of Yes-associated protein (Yap) and its cofactors, TEAD family members (Zhao et al., 2008). CTGF activates ischemic heart disease cardiomyopathy in both mice and humans by activating the Yap1/Taz/Tead1 pathway (Hou et al., 2017).

The transcription factor TEAD (TEA/ATTS domain) is a conserved nuclear protein and plays a pivotal role during lineage development, including trophoectoderm specification, cardiogenesis and myogenesis (Chen et al., 1994; Yagi et al., 2007). Teads activate Yap1 which stimulates cardiomyocyte proliferation during embryonic mouse heart development (von Gise et al., 2012; Xin et al., 2011). It has also been shown that disruption of *Tead1* in the embryonic mouse heart results in heart defects and embryonic lethality (Chen et al., 1994). Moreover, overexpression of Tead1 in post-natal mouse heart results in contractile dysfunction of cardiomyocytes and cardiac fibrosis, hallmarks of heart disease (Tsika et al., 2010). In addition, transgenic mice with cardiac muscle-specific over-expression of Tead4 have been shown to increase atrial weight and exhibit defects in cardiac conduction (Chen et al., 2004). Furthermore, a mutual regulation of Tead4 and CTGF requires trophectoderm specification for proper cellular differentiation in preimplantation embryo development (Akizawa et al., 2019).

Here we investigated the function of a mouse ESC-specific lncRNA, *Platr4* (pluripotency-associated transcript 4). We found that CRISPR/Cas9-mediated deletion of *Platr4* in ESCs disrupts mesoderm/endoderm lineage specification, specifically the cardiac lineage, while preserving self-renewal and pluripotency. We further show that *Platr4* is necessary for the activation of the core cardiac gene expression network that induces cardiac transcription factors (Mesp1, Gata4, Tbx5, Nkx2.5) and initiates EMT by inducing *N-Cadherin* and reducing *E-Cadherin* expression. Further analysis revealed that *Platr4* regulates genes at the transcriptional level, and the loss of *Platr4* in ESCs results in a significant depletion of Ctgf at both the RNA and protein levels. Moreover, over-expression of *Ctgf* rescued the *Platr4* knockout phenotype, demonstrating that *Platr4* functions in *trans* upstream of *Ctgf*. To carry out its function *Platr4* interacts with the transcription factor, Tead4 and together they modulate their downstream target, *Ctgf*, for cardiac cell-fate specification. Importantly, deletion of *Platr4* in a knockout mouse model results in a pathophysiological condition of myocardial atrophy, valve mucinious degeneration associated with decreased cardiac output and myocyte contractility. Together, our findings reveal a novel mechanism of action of the *Platr4* lncRNA-based complex where the *Platr4*/Tead4/*Ctgf* axis plays an essential role in cardiac lineage commitment.

## Results

We identified *Platr4* lncRNA to be specifically expressed in mouse ESCs in a differential RNA-seq screen upon differentiation of ESCs to neural progenitor cells (NPCs) (Bergmann et al., 2015). Rapid amplification of cDNA ends (RACE) cloning followed by Sanger sequencing indicated *Platr4* to be a 1,034 nucleotide-long polyadenylated RNA encoded from two exons, consistent with our RNA-seq and northern blot analysis (Figure 1A, 1B and S1A). Examination of the *Platr4* sequence with the Coding Potential Calculator (CPC) (Kong et al., 2007), Coding-Potential Assessment Tool (CPAT) (Wang et al., 2013), and PhyloCSF (Lin et al., 2011) indicated that the *Platr4* transcript does not have protein coding capacity (Figure S1B and S1C). Publicly available ENCODE RNA-seq datasets of adult mouse tissues showed that *Platr4* is expressed in ESCs but not in NPCs or any adult tissues (Figure 1C). Cellular fractionation of ESCs followed by quantitative RT-PCR (qRT-PCR) demonstrated that *Platr4* is enriched in the nucleus, and especially in the chromatin fraction (Figure 1D). Consistent with the nuclear fractionation, single-molecule RNA fluorescent in situ hybridization (smRNA-FISH) revealed that *Platr4* is predominantly localized in the nuclei (Figure 1E). In addition, we performed smRNA-FISH on whole mount and paraffin sections of embryos ranging from embryonic day 3.5 (E3.5) to 12.5 (E12.5). We found prominent expression of *Platr4* in both trophectoderm and the inner cell mass of E3.5 embryos (blastocysts) (Figure 1F), consistent with qRT-PCR data in preimplantation embryos (Figure S1D). We also detected prominent expression of *Platr4* in extra embryonic ectoderm at E6.5 (Figure S1E) which is derived from the trophectoderm lineage and contributes to placenta development. Notably, we detected very low expression of *Platr4* at E8.5 and E10 (Figure S1E) and no expression observed from E12.5 (Figure S1E) and later stage embryos (data not shown), suggesting that *Platr4* is only expressed during early embryonic development. Thus, *Platr4* is a nuclear-enriched lncRNA that is expressed in ESCs and in early development.

**Figure 1:**
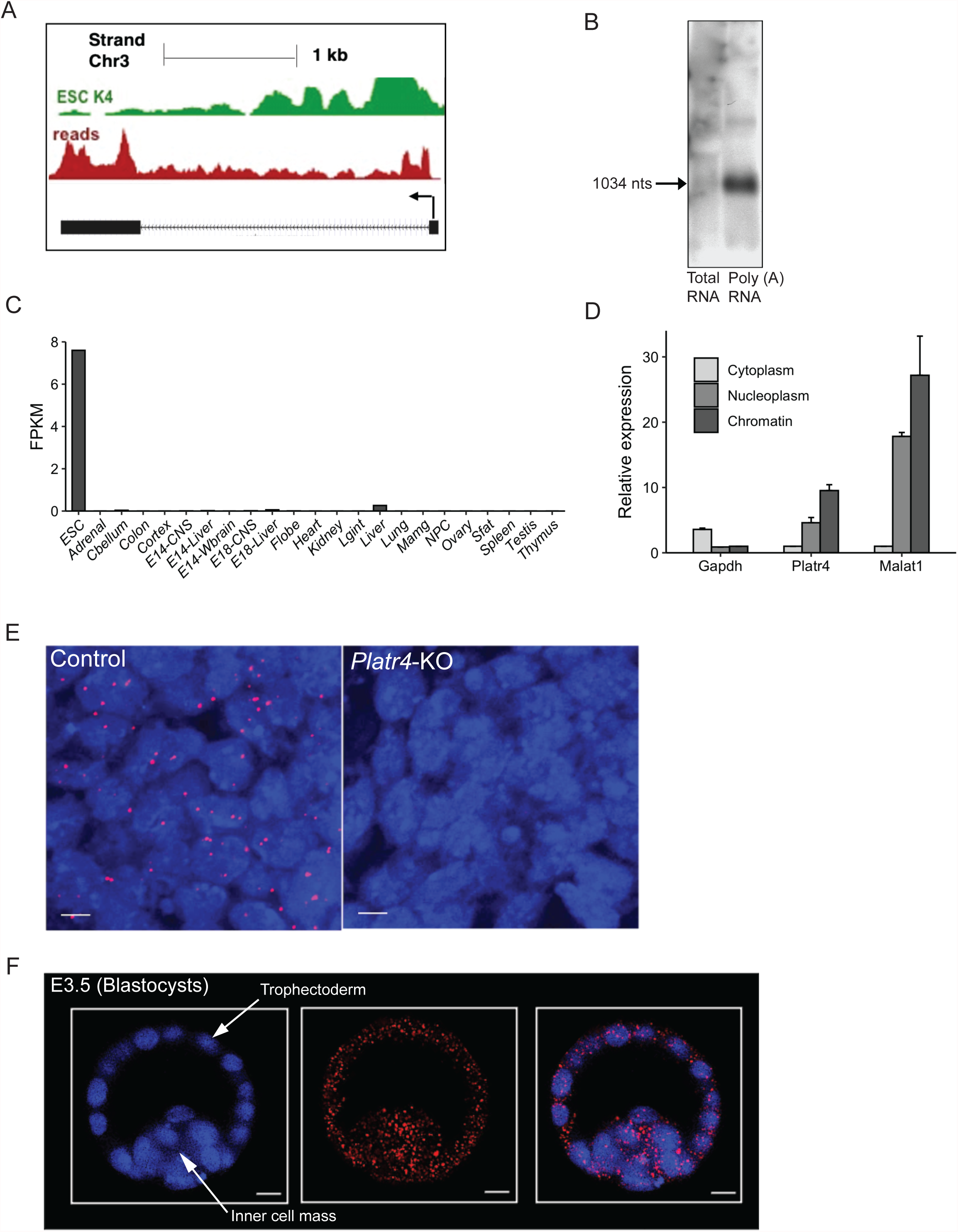
Platr4 is an ESC-specific nuclear enriched lncRNA. (A) UCSC genome browser view of the *Platr4* locus. Also shown are ChIP-seq tracks for H3K4me3 and RNA-seq reads of ESCs. (B) Northern blot data showed *Platr4* is 1,034 nucleotides in length (C) Fragments per kilobase per million mapped reads (FPKM) values for *Platr4* in various mouse tissue types from ENCODE data sets. (D) Subcellular fractionation followed by qRT-PCR showed the localization of *Platr4* transcripts. *Gapdh* and *Malat1* were used as cytoplasmic and nuclear markers for quality control. Data are presented as mean values ± SD (n = 3 independent experiments). (E) Single-molecule RNA-FISH images indicate localization of *Platr4* RNA transcripts (red dots) within nuclei in wild-type ESCs. No detectable *Platr4* transcripts are observed in *Platr4*-KO ESCs. Scale bars are 50 μm. (F) Single-molecule RNA-FISH in pre-implanted embryos (blastocysts, E3.5). Scale bars are 50 μm.

To dissect the molecular function of *Platr4*, we developed a genetic knockout (KO) of *Platr4* in mouse embryonic stem cell (mESC) lines (V6.5 and AB2.2) using the CRISPR/Cas9 approach. We designed two sgRNAs spanning the promoter region (+300 bp / −200 bp) relative to the transcription start site (TSS) of *Platr4* to create a genomic deletion verified by genomic PCR and Sanger sequencing (Figure 2A and S2A). We performed nucleofection of transiently expressing Cas9 and guide RNAs in order to enhance Cas9 specificity and reduce off-target activity (Slaymaker et al., 2016). Furthermore, transfection efficiency (40-50%) in transient nucleofection is higher than viral transduction (1%) in mESCs. Guide RNA targeting Renilla luciferase was used as a non-targeting control. Luciferase control vs *Platr4*-KO cells were single-cell sorted 48 hours after nucleofection. We detected a complete loss (99%) of *Platr4* transcript in ESCs and verified by northern blot, qRT-PCR, and as well as smRNA-FISH (Figure 2B, 1E and S2B). Interestingly, the loss of *Platr4* in ESCs did not affect colony morphology, cell cycle kinetics (data not shown), proliferation (Figure S2C) or expression of master pluripotency factors, such as Pou5f1 (Oct4) and Nanog (Figure 2C and S2D). Thus, *Platr4* is not essential to maintain ESC pluripotency and self-renewal.

**Figure 2:**
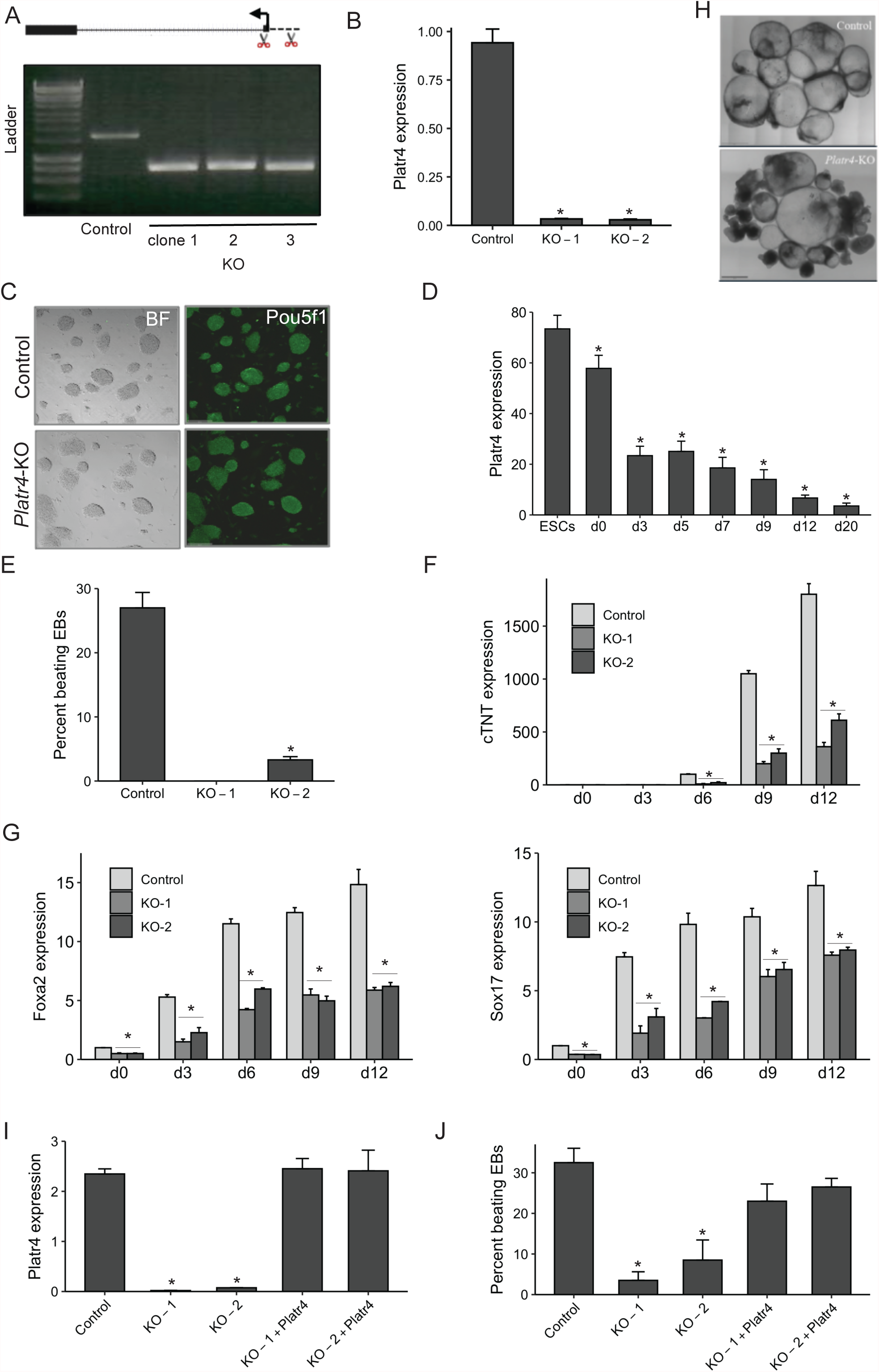
Platr4 is essential for ESC differentiation. (A) CRISPR/Cas9-mediated deletion of *Platr4*-KO clones in ESCs. A pair of sgRNAs near the transcription start site of *Platr4* were used to establish a ∼600bp genomic deletion. Knockout clones were verified by genomic PCR (clones 1, 2, 3), and a Renilla Luciferase sgRNA was used as a negative control. (B) Deletion of *Platr4* transcripts was further verified by qRT-PCR. (C) Deletion of *Platr4* in ESCs did not affect colony morphology and immunostaining of Pou5f1. Scale bar, 150 μm. BF stands for bright field. (D) *Platr4* level in differentiated embryoid bodies at different days. (E) Loss of percentage of spontaneously beating EBs in *Platr4*-KO clones compared to control at day 12 of differentiation n= 100 EBs was counted, and experiments were performed in triplicate. (F) The relative level of cardiac Troponin T (cTnT) was measured by qRT-PCR using control vs. *Platr4*-KO EBs at different time points upon differentiation. (G) qRT-PCR analysis of reduced expression levels of endoderm-specific germ layer markers (*Foxa2* and *Sox17*) in *Platr4*-KO EBs compared to control. (H) Control vs. *Platr4*-KO EBs at day 12. Platr4-KO EBs show a smaller size and darker cavity compared to control EBs. (I) Ectopic expression of *Platr4* in *Platr4*-KO ESCs measured by qRT-PCR. (J) Ectopic expression of *Platr4* in *Platr4*-deleted EBs can rescue the percentage of contracting EBs. Data are presented as mean values ± SD. All experiments were performed in triplicate. *p < 0.05 (student’s t-test).

ESCs have the capacity to differentiate into derivatives of the three germ layers: endoderm, mesoderm and ectoderm. Given its expression in early embryos, we choose to study the impact of the loss of *Platr4* in lineage commitment and differentiation. We next induced control vs. *Platr4*-KO ESCs to differentiate through withdrawal of leukemia inhibitory factor (LIF) by allowing cells to aggregate into embryoid bodies (EBs) by the hanging drop method. The EBs form cell types corresponding to all three germ layers and are a routinely used model system to assess the early differentiation potential of ESCs (Desbaillets et al., 2000). We further confirmed that *Platr4* lncRNA is expressed in ESCs and is significantly down-regulated upon their differentiation (Figure 2D). During the differentiation process, we first observed abnormalities in the spontaneous contraction of EBs in *Platr4*-depleted cells. In control cells, 27% of EBs exhibit beating by day 12 compared to 0% (clone-1) and 3% (clone-2) in *Platr4* KO cells (Figure 2E). Consistent with this observation, the expression of cardiac Troponin T (cTnT) and myosin heavy chain 7b (myh7b), two important proteins involved in cardiomyocyte contraction, was evaluated by qRT-PCR at different time points and showed decreased levels in *Platr4*-depleted EBs (Figure 2F and S2E). We also found that *Platr4*-depleted EBs exhibited significantly decreased levels of the definitive endoderm (DE) markers, *Sox17* and *Foxa2* (Figure 2G) without affecting the neuroectoderm markers (Figure S2F). In addition, morphological abnormalities of EBs were observed with smaller size and darker cavities in *Platr4*-depleted compared to control EBs (Figure 2H), consistent with the observation that bright cavity EBs have better differentiation capacity than dark cavity EBs (Kim et al., 2011). Examination of histological sections of *Platr4*-KO EBs at day 12 of differentiation showed a range of tissues, such as neural rosettes, gut endoderm, and cartilage (Figure S2G), suggesting that *Platr4* is likely not required for global ESC differentiation. To exclude the possibility that the phenotypes observed in *Platr4*-KO cells were caused by disturbing chromatin structure rather than specific loss of the *Platr4* transcript, we generated single-cell ectopic expression rescue clones of *Platr4* in *Platr4*-depleted ESCs. Notably, transient rescue of *Platr4* significantly increased the percentage of beating EBs (Figure 2I and 2J), indicating that the *Platr4* transcript plays an important role in these processes *in situ*, and likely exhibits its effect in *trans*. Together, these data demonstrate that *Platr4* may function in mesoderm and endoderm cell fate specification.

In order to examine whether *Platr4* functions in cis or in trans we examined the expression levels of neighboring genes (*Sclt1, Jade1, D3Ertd751e*) within a 100kb window and determined an equivalent level of expression in *Platr4*-KO and control cells (Figure S3A), consistent with a role for *Platr4* exerting its function in *trans*. LncRNAs play an important role in the molecular circuitry of the undifferentiated ESC state to control pluripotency and lineage differentiation (Guttman et al., 2011). Therefore, to explore the transcriptional control of *Platr4* in ESC state in order to understand its function during differentiation, we analyzed global gene expression profiles using poly(A)+ RNA-seq on control vs. *Platr4*-KO ESCs. To this end, a large number of differentially expressed genes (DEGs) (scaled expression) were identified between *Platr4*-KO and control ESCs (Figure 3A, 3B, S3B and Supplementary data 2). The top ten significantly down-regulated genes were validated by qRT-PCR (Figure 3C). DAVID GO term pathway analysis in control vs. *Platr4*-KO ESCs showed that DEGs was significantly enriched into eight significant pathways, including ECM-receptor interaction, focal adhesion, and PI3K-Akt signaling pathway, all of which play a vital role in mammalian development (Chatzizacharias et al., 2010; Riley et al., 2005) (Figure 3D). Notably, customized GSEA analysis (Fagnocchi et al., 2016) using DEGs revealed significant upregulation in pluripotency (Figure S3C and Supplementary data 3) although no change was observed in the master pluripotency transcription factors Pou5f1 and Nanog (Figure 2C and S2C). We further performed an RNA-seq experiment using control vs *Platr4*-KO EBs and DEGs (log fold change) were analyzed at different days (D0-D12) (Figure 3E and Supplementary data 4). DAVID GO term analysis showed DEGs were significantly enriched in eight pathways, including ECM-receptor interaction, focal adhesion, PI3K-Akt signaling pathway, pathways in cancer, proteoglycans in cancer, endocytosis, coagulation cascade, and amoebiasis (Figure S3D). Next, customized GSEA analysis showed that DEGs were significantly down-regulated in the mesoderm and endoderm lineages without altering the neuroectoderm lineage (Figure 3F and Supplementary data 3), consistent with our previous observation that *Platr4* is critical to mesoderm and endoderm lineage specification (Figure 2F and 2G). These results indicate that *Platr4* functions in *trans* and interacts with transcription factors (TFs) to regulate the expression level of target genes in ESC state. Therefore, we performed iRegulon (Cytoscape) analysis (Janky et al., 2014) to predict the potential TFs from the co-expressed gene set of *Platr4* lncRNA. We used a Normalized Enrichment Score (NES) >3 for the significant enrichment of target TFs. iRegulon interactome maps with TFs using up-regulated and down-regulated DEGs were found. Transcription factors, Zfp143, E2f, and Tead family were enriched with *Platr4* co-expressed genes in up-regulated and down-regulated DEGs (Figure 3G and Supplementary data 5).

**Figure 3:**
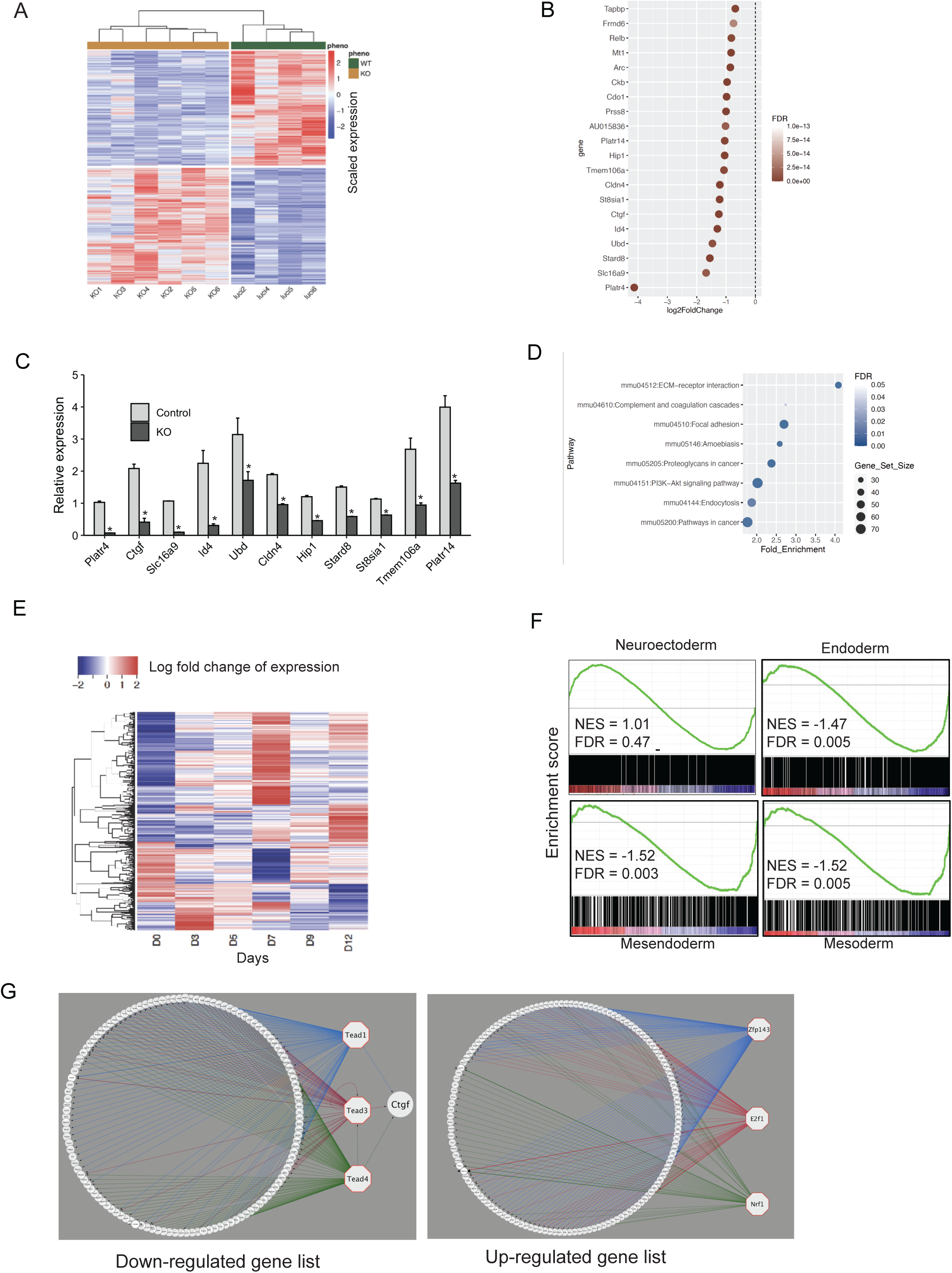
Platr4 regulates mesoderm/endoderm lineage specification. (A) Heat-map of differently expressed genes (DEG) (scaled expression) in luciferase control and *Platr4*-KO ESCs. (B) Top 20 downregulated genes upon deletion of *Platr4* in ESCs. (C) The top ten downregulated genes were quantified by qPCR using control vs. downregulated genes in KO ESCs. Data are presented as mean values ± SD. (n = 3 independent experiments). (D) Analysis of significant GO terms in downregulated DEG that were dysregulated upon deletion of *Platr4* in ESCs. (E) Heat-map of DEG (log fold change) in luciferase control and *Platr4*-KO EBs upon differentiation at various days. (F) Gene set enrichment analysis profiles of DE genes. The gene lists are shown in Table S1. (G) iRegulon analyses detected the enriched transcription factor motif in downregulated (left panel) and upregulated (right panel) DE genes visualized by Cytoscape software. Normalized Enrichment Score (NES) >3.

iRegulon analysis revealed that *Platr4* might exert its function via the regulation of transcription factors of the Tead family and its known downstream target genes, *Ctgf* (Liu et al., 2016a; Zhao et al., 2008). First, we found that the expression levels of both RNA and protein of Tead1, 2, 3, 4 were equivalent in control and *Platr4*-KO ESCs (Figure 4A, 4B and S4A). Next, we performed siRNA knockdown analysis of each Tead family member and found that they did not affect *Platr4* expression level (Figure 4C and S4B), although the immediate downstream targets of Tead4 (*Ctgf, Gli2, Vgll3*) (Zhou et al., 2016) were significantly down-regulated (Figure 4C and 4D). Interestingly, we found that *Ctgf* is one of the top five significantly down-regulated gene in RNA-seq data (Figure 3B and 3C). Furthermore, we found that both the RNA and protein levels of Ctgf were significantly reduced in *Platr4*-KO vs control ESCs (Figure 4E, 4F-upper and lower panel). Moreover, ectopic expression of *Platr4* in *Platr4*-KO ESC resulted in a corresponding increase in the mRNA level of *Ctgf* (Figure 4G), but not the level of *Gli2* or *Vgll3* (Figure S4C). These results suggest that *Ctgf* is a direct downstream target of *Platr4*, and it functions in *trans*. We next performed CRISPR/Cas9 knockdown (KD) using gRNAs targeting *Ctgf* in ESC puromycin-resistant cells for downstream functional assays. Both the RNA and protein expression levels of Ctgf in *Ctgf*-KD ESC were verified (Figure 4H and 4I). Control and *Ctgf*-depleted ESCs were induced to differentiate via withdrawal of LIF by allowing cells to aggregate into EBs by the hanging drop method. We found that the reduction of *Ctgf* in EBs at day12 phenocopied the *Platr4*-KO EBs and exhibited a significant reduction in spontaneous contracting EBs: 24% of EBs showed an observable beat at day 12 in control cells compared to 14% in *Ctgf*-KD cells (Figure 4J). Notably, ectopic expression of the full-length Ctgf gene in *Platr4*-KO ESC increases the percentage of contracting EBs in *Platr4* depleted cells (Figure 4K). Together, these data suggest that *Ctgf* is a potential regulator of cardiomyocyte differentiation and a critical downstream target of *Platr4*.

**Figure 4:**
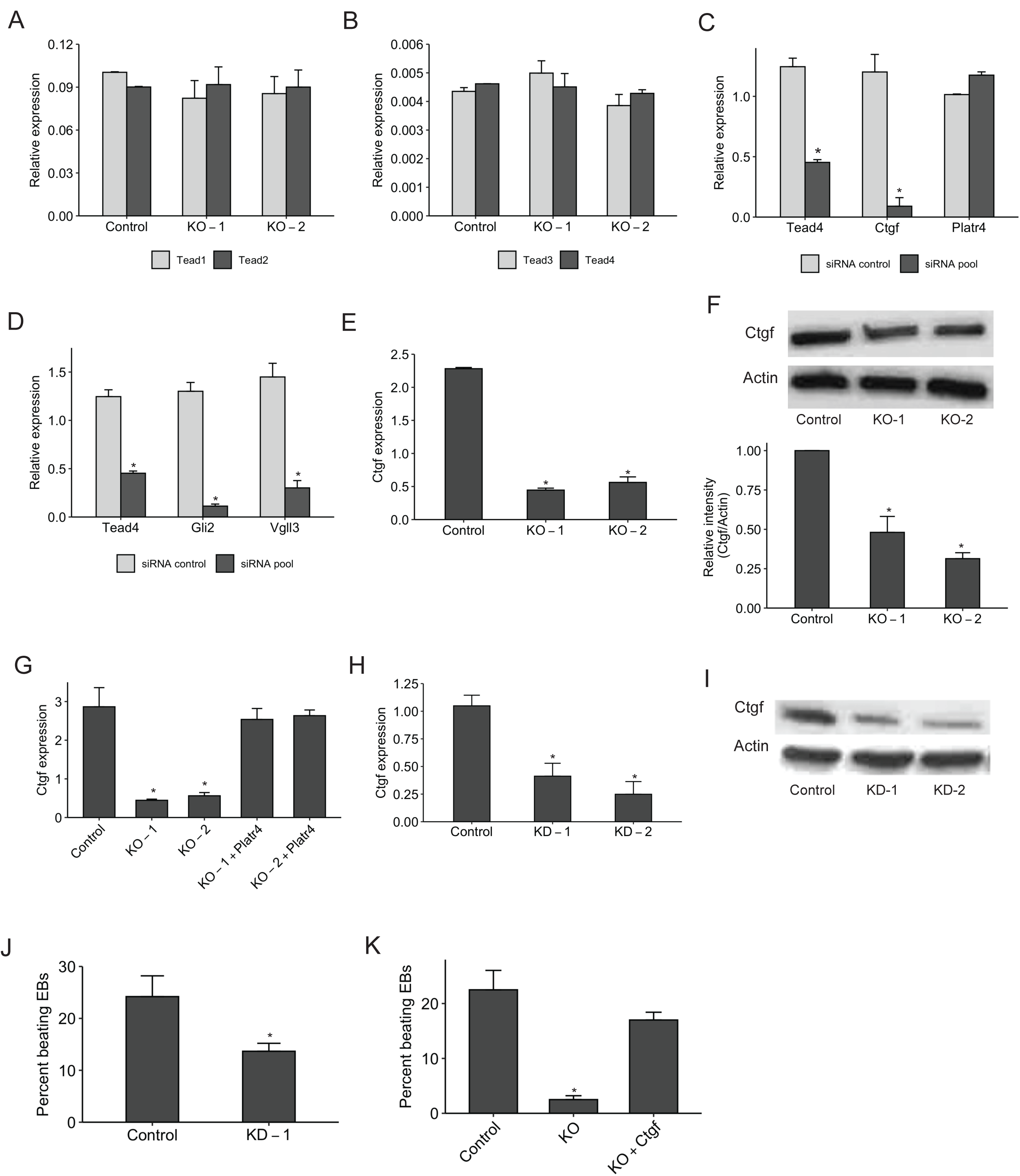
Platr4 functions upstream of connective tissue growth factor (Ctgf) (A & B) The relative expression level of *Tead1, Tead2, Tead3, Tead4* in control vs. *Platr4*-KO ESCs, measured by qRT-PCR. (C & D) qRT-PCR analysis of Tead4-downstream target genes (*Ctgf, Gli2, Vgll3*) and *Platr4* in ESCs using Tead4-siRNA and control siRNA. (E) Relative expression of *Ctgf* in *Platr4*-KO vs. control ESCs. (F) Western blot analysis of Ctgf level (upper and lower panel) in control and *Platr4*-KO ESCs (n=2 independent experiment). (G) *Ctgf* level is rescued upon ectopic expression of *Platr4* in *Platr4*-KO cells as determined by qRT-PCR. (H & I) CRISPR/Cas9-mediated knockdown of *Ctgf* in ESCs, verified by both qRT-PCR and western blot analysis (n=2 independent experiment). KD stands for the knockdown. (J) Reduced percentage of contracting EBs upon downregulation of Ctgf compared to control. (H) Ectopic expression of *Ctgf* in *Platr4*-KO cells rescues the percentage of contracting EBs in *Platr4*-KO ESCS upon differentiation. All experiments were performed in triplicate. Results are mean ± SD (n=3) *p < 0.05 (student’s t-test).

Next, a directed differentiation technique was applied to recapitulate aspects of normal early cardiac development. Differentiation of ESCs into cardiomyocytes (CM) from the mesoderm lineage are assessed by the initial expression of Brachyury (T) and Eomesodermin (Eomes) (Klattenhoff et al., 2013; Murry and Keller, 2008). Therefore, to study *Platr4* function in cardiac cell fate from the mesodermal lineage, we employed a directed *in vitro* CM differentiation assay that permits isolation of cell populations at well-defined stages [ESC, EB, MES (precardiac mesoderm), CP (cardiac progenitors, and CM] (Klattenhoff et al., 2013) (Figure 5A). Using this assay, we found that *Platr4*-KO EBs were smaller in size and exhibited an irregular shape at day 4 MES compared to control (Figure 5B), although no morphological changes were observed at day 2, a time point that precedes the addition of cardiac growth factors. Despite, the relative elevated levels of *T* and *Eomes* in MES and CP populations in *Platr4*-KO compared to control cells (Figure 5C), expression of core cardiac transcription factors at day 4 MES, day 6 CPC, and day12 CM were significantly downregulated in *Platr4*-KO cells (Figure 5D, 5E, and 5F). Interestingly, cell surface markers, *Flk1* and *PdgfRα* that are enriched in CP populations were also down-regulated in *Platr4*-KO cells (Figure 5G). *Flk1* and *PdgfRα* represent early cardiac mesoderm markers (Liu et al., 2016b). These data further support that loss of *Platr4* affects the cardiac lineage.

**Figure 5:**
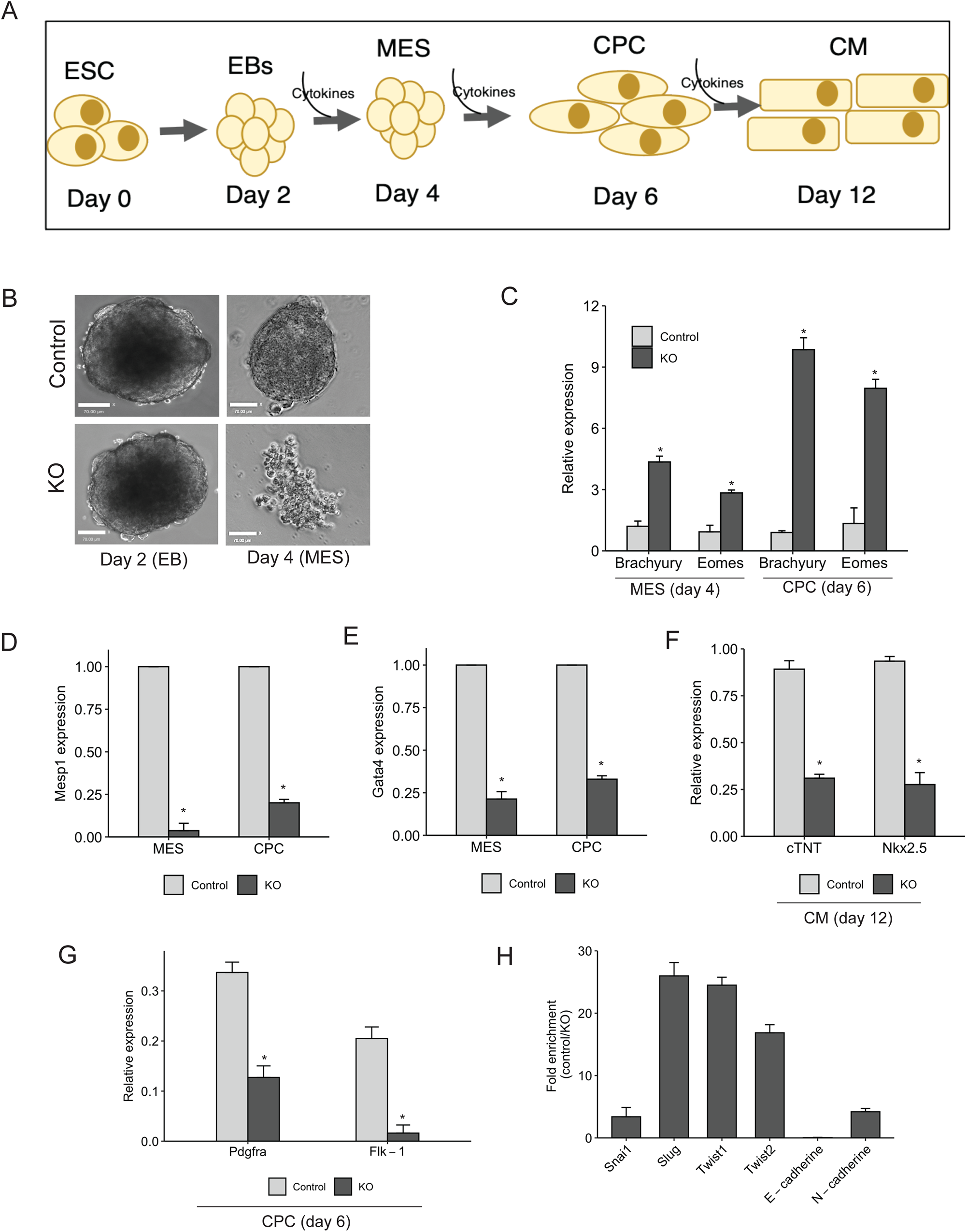
Directed cardiac differentiation. (A) ESCs were differentiated into cardiomyocytes (CM) and progressed through mesoderm (MES) and cardiac progenitor cell (CPC) using serum free media with the sequential addition of cytokines. (B) *Platr4*-KO EBs showed a smaller size at the MES stage compared to controls at day 4. Scale bar, 50 μm. (C) The relative expression level of *Brachyury* and *Eomes* were elevated compared to control at respective days. (D, E, and F) Relative expression of core cardiac transcription factors displayed significant downregulation in *Platr4*-KO compared to control cells upon different stages of differentiation. (G) The relative level of cell surface markers (*Pdgfrα* and *Flk-1*) was reduced in *Platr4*-depleted CPCs. (H) qRT-PCR measurement of the altered expression levels of EMT genes in *Platr4*-depleted CPCs. All experiments were performed in triplicate. Values are mean ± SD (n=3) *p < 0.05 (two-tailed student’s t-test).

Cell-ECM interactions are crucial for cardiac lineage differentiation (Rozario and DeSimone, 2010; Zhang et al., 2012). We found that depletion of *Platr4* in ESCs significantly reduced ECM genes compared to control cells (Figure S5). Interestingly, the Snail-Ctgf axis induces epithelial-to-mesenchymal transition (EMT) to differentiate fibroblasts into myofibroblasts (Lee et al., 2013), suggesting that Ctgf regulates EMT genes. We examined the expression of EMT genes such as Snai1 and Slug (Aban et al., 2021; Lim and Thiery, 2012), as well as Twist1 and Twist2 (Yang et al., 2004) and found that they were induced in the CP population (Figure 5H). We found that deletion of *Platr4* reduces the expression of *E-Cadherin* and induces N-Cadherin expression (Figure 5H) consistent with the initiation of EMT (Aban et al., 2021; Lim and Thiery, 2012). These result highlight the importance of *Platr4* in the initiation of EMT and indicate that *Platr4* is required in cardiac cell specification at a critical point in the mesoderm lineage.

Ctgf is a direct downstream target of the Tead transcription factor (Zhao et al., 2008). Since we found significant enrichment of the Tead family in downregulated *Platr4*-DEGs (Figure 3G) we performed RNA immunoprecipitation (RIP) using antibodies against Tead1, 2, and 4 to assess specific interactions with *Platr4*. Interestingly, we found a specific interaction of *Platr4* with Tead4 (Figure 6A-upper and lower panel) but none with Tead1,Tead2 and Tead3 (Figure S6A, B and C). Next, using *in vitro* biotin-RNA pull-down, we further confirmed this specific interaction of Tead4-*Platr4* in ESCs, but not with Tead1 and Tead2 (Figure 6B). In addition, to examine the regulatory role of Tead4 on Ctgf in ESCs, we performed either ectopic over-expression (Figure 6C and 6D) or siRNA-mediated down-regulation of Tead4 (Figure 4C) to demonstrate the up-regulation or down-regulation of *Ctgf*, respectively. These results verify that modulation of Tead4 expression level influences the expression level of Ctgf in ESCs, suggesting that the *Platr4*/Tead4 RNP complex is essential for regulating *Ctgf*. We further performed chromatin immunoprecipitation (ChIP) using a Tead4 antibody, and multiple qPCR primer pairs showed that Tead4 has a high occupancy over the *Ctgf* promoter, as also suggested in other studies (Benhaddou et al., 2012; Kang et al., 2018). Notably, the occupancy was impaired in *Plar4*-KO ESCs and was able to be restored upon ectopic expression of *Platr4* in *Platr4*-KO ESCs (Figure 6E). Together, these results provide compelling evidence that the interaction of Tead4 protein with *Platr4* lncRNA is required for Tead4 to bind to regulatory motifs in the *Ctgf* gene in ESCs. This is the first demonstration that the interaction between a lncRNA and Tead4 are necessary to modulate a down-stream target gene and regulate cardiac lineage differentiation.

**Figure 6:**
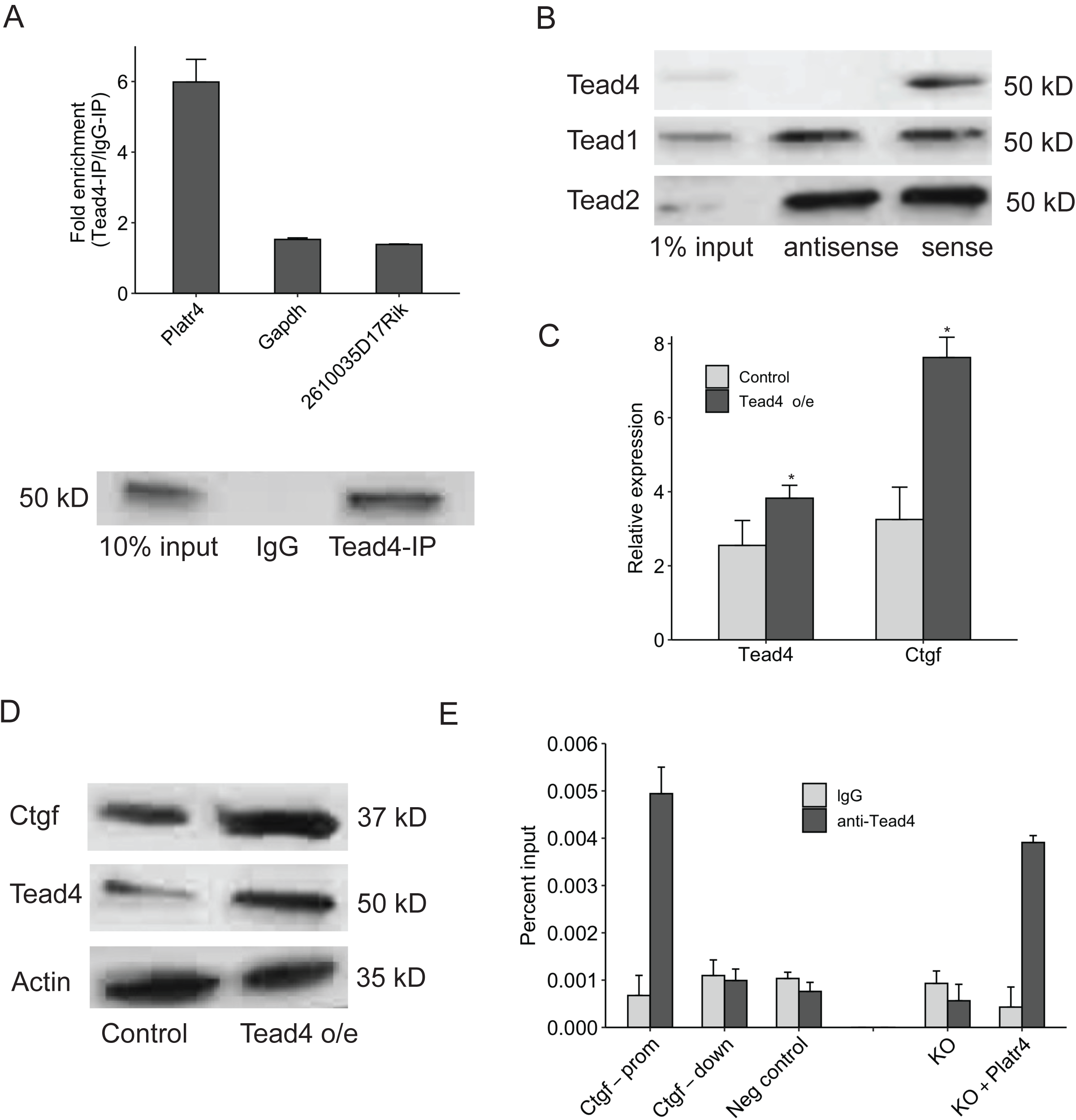
Platr4 interacts with Tead4. (A) RIP assay confirmed the interaction of *Platr4* with Tead4 using Tead4 antibody. Fold enrichment of *Platr4* over IgG signal is shown. *Gapdh*, and *2610035D17Rik* (lncRNA) transcripts were used as controls (upper panel). Western blot analysis of Tead4 was performed to confirm downregulation (lower panel). (B) Biotin-RNA pull-downs using full-length *Platr4* transcript in nuclear ESC extracts showed specific binding to Tead4 but not with Tead1 and Tead2. (C) qRT-PCR analysis of increased *Ctgf* expression in ectopic over-expression of Tead4. (D) Western blot analysis of Ctgf expression in ectopic over-expression of Tead4. (E) ChIP-qPCR analysis displayed Tead4 occupancy over *Platr4* targeting region of the *Ctgf* DNA locus in control, *Platr4*-KO, and ectopic *Platr4* expression in *Platr4*-KO ESCs. *Ctgf*-prom, *Ctgf*-down stands for promoter and downstream gene. *Gapdh* was used as a negative control. All experiments were performed in triplicate. Values are mean ± SD (n=3) *p < 0.05 (two-tailed student’s t-test).

We generated a *Platr4*-KO mouse in the C57BL/6J strain using two sgRNAs targeting the TSS and first exon of *Platr4* as described earlier (Figure 2A), and three KO founders were genotyped (Figure S7A) and used to establish three independent lines. The *Platr4*-KO alleles were backcrossed to C57/BL6 background for ten generations to yield a pure C57/BL6 genetic background. Heterozygotes intercrossed with WT mice generated wild-type and *Platr4*-KO littermates (Figure 7A and 7B). All homozygous KO mice used for breeding were fertile, and they did not show any gross physical abnormalities compared to WT mice. Interestingly, we observed sudden death of young adult and adult mice (40%) in the KO cohort but not in the WT cohort (data not shown). When we examined the hearts of the young adult dead mice (N=3) and live mice (N= 10 for both WT and KO cohorts) by H&E staining of tissue sections (Figure S7B), we found 60% of the knockout mice exhibited valve defects with fibrocartilaginous metaplasia (Figure 7C and S7C), fibro osseous metaplasia and mucinous degeneration compared to WT (Figure 7D and S7D) (N=10). In addition, we have found perivascular and myocardial mineralization (60% KO-mice, (N=10/group) (Figure 7E) and myocardial atrophy and fibrosis (S7E). To further assess heart defects, we performed echocardiography analysis using WT vs. KO mice. Echocardiography is used to visualize the cardiovascular structures and measure cardiac function in mice due to its advanced spatial-temporal resolution. Our findings showed a significant 30% decrease in the percentage of fractional shortening (% FS) without altering the percentage ejection fraction (EF%) in KO compared to WT mice (Figure 7F). This observation supports our *in vitro* data of cardiomyocyte dysfunction since percent FS indicates changes in left ventricle (LV) chamber size and myocyte contractility (Gardin et al., 1995). We further demonstrated increased ventricular wall thickness and ventricular mass in KO vs WT mice indicating ventricular hypertrophy (Figure 7G and 7H). Interestingly, significant reduction of cardiac output (CO) and heart rate (HR) in KO compared to WT mice (Figure 7I) may explain our observation of heart failure and sudden death since CO is an important measuring parameter of cardiac dysfunction (Hoffman et al., 2019). Thus, genetic loss of *Platr4* impacts both *in vitro* as well as *in vivo* cardiac development.

**Figure 7:**
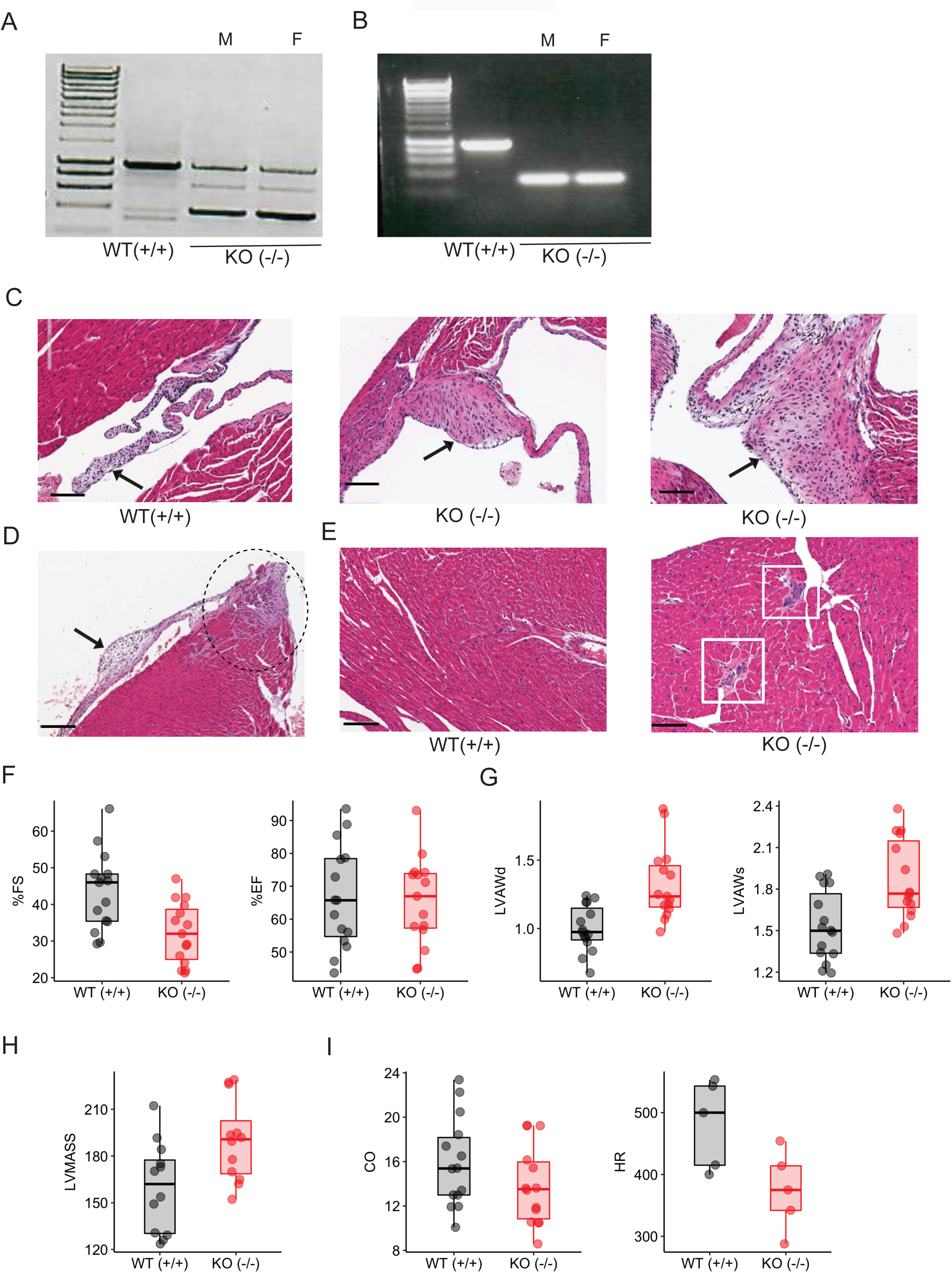
Phenotype of Platr4-KO mice. (A & B) PCR genotyping for *Platr4* WT (+/+), homozygous (-/-), and heterozygous (+/-) KO mice. M and F stand for male and female. (C) H&E staining (longitudinal sections) of the adult heart from WT mice showed normal valve, and *Platr4*-KO mice showed valve defects with fibro cartilaginous metaplasia. Scale bar: 100 μm. (D) H&E staining for the adult *Platr4*-KO heart showed fibrocartilaginous metaplasia and myxomatous degeneration of the valves. Scale bar: 100 μm. (E) H & E staining of KO mice showed mineralization, but the WT heart was normal. Scale bar: 100 μm. (F) Echocardiography analysis for the %FS and %EF of WT and *Platr4*-KO mice. (G) Echocardiography analysis for LVAW in diastole or systole in WT and *Platr4*-KO mice. (H & I) Echocardiography analysis of LV mass, cardiac output (CO), heart rate (HR) (n = 5) in WT and *Platr4*-KO mice. FS= fraction shortening, EF= ejection fraction, LVAWd= left ventricular anterior wall thickness in diastole, LVAWs= left anterior ventricular wall at end systole, LV= left ventricle. Values are mean ± SEM (n=15/group) *p < 0.05 (two-tailed student’s t-test).

## Discussion

Dynamic regulation of transcriptional programs is vital in lineage specification during mammalian development, and disruption or faulty regulation of these processes can lead to various developmental disorders. *Bvht* was the first identified lncRNA in mice to display a crucial role in cardiac commitment, however, its loss had no phenotype in a knockout mouse model and a human ortholog has thus far not been identified (Han et al., 2018; Klattenhoff et al., 2013). More recent studies have shown the epigenetic regulation of lncRNAs in mesoderm and endoderm lineage commitment and their role in embryonic heart development (Grote et al., 2013; Guo et al., 2018). Recently, the muscle-specific lncRNA *Charme* has been reported to control skeletal and cardiac myogenesis in *in vitro* and *in vivo* systems (Ballarino et al., 2018). In addition, the chromatin-associated lncRNA *Myheart* was shown to be highly expressed in the adult mammalian heart and it has been shown to repress cardiomyopathy in the mouse heart (Han et al., 2014).

Here we report for the first time on the mechanism by which a nuclear enriched lncRNA *Platr4*, exclusively expressed in ESCs and whose loss significantly represses downstream endoderm differentiation and cardiac lineage differentiation in both EBs and directed CM differentiation, yet does not impact pluripotency. Interestingly, *Platr4*-KO mice showed valve defects, myocardial atrophy and cardiac dysfunction, suggesting that deletion of *Platr4* in mice impairs heart function. Together, our findings reveal that *Platr4* is required for proper cell fate specification and thus serves as an *in vivo* functional lncRNA in murine mesoderm/endoderm lineage commitment.

We identified the involvement of *Ctgf* in cardiomyocyte fate determination and function as a downstream target of *Platr4* lncRNA. *Ctgf* is essential for embryonic development, and global *Ctgf*-KO mice displayed skeletal anomalies and perinatal death (Lambi et al., 2012). Abnormal Ctgf level is associated with multiple disorders, including fibrosis and cancer in many organs and tissues (Ramazani et al., 2018). A significant pathological condition of heart disease is cardiac fibrosis due to excessive deposition of extracellular matrix components in the heart. CTGF regulates the fibrotic process as a matrix protein and is a known biomarker for cardiac fibrosis (Daniels et al., 2009). It has also been shown to be required for initiating the fibrosis process of several organs and tissues, including the heart (Shi-Wen et al., 2008). We also show the cardiac fibrosis in the *Platr4*-KO mice. Moreover, plasma CTGF concentration is used as a biomarker for cardiac dysfunction in chronic heart failure patients (Koitabashi et al., 2008). Here, we report for the first time that depletion of Ctgf in ESCs impacts cardiomyocytes contractility during lineage differentiation, consistent with the finding that the Ctgf protein is involved in cardiomyocyte differentiation (Han et al., 2014). In addition, cell-ECM interactions are critical for gastrulation and morphogenic patterning in mammalian development, including normal cardiac development (Gustafsson and Fassler, 2000; Kleinman et al., 2003; Lockhart et al., 2011). ECM components are also critical regulators for *in vitro* differentiation of ESCs and somatic cell reprogramming at different developmental stages (Goh et al., 2013; Sart et al., 2014). CTGF is a cysteine-rich, ECM-associated protein and a direct downstream modulator of the TGF-β signaling pathway during heart development (Chen et al., 2000; Qi et al., 2007; Song et al., 2007). We found that depletion of *Platr4* affects ECM components and downregulation of the *TGF-β* gene (data not shown). Moreover, Ctgf regulates EMT genes to differentiate fibroblasts into myofibroblasts (Lee et al., 2013). Our findings show that *Platr4* is required to initiate the EMT gene network. This suggests that *Platr4* may be involved in cardiovascular lineage differentiation through the structural support network of functional tissue.

The mammalian Tead/Tef family consists of four members, Tead1-4 with similar domain architecture, and each Tead has a tissue-specific expression (Anbanandam et al., 2006; Jacquemin et al., 1996; Jacquemin et al., 1997). Tead4 plays an essential role in trophectoderm lineage determination (Nishioka et al., 2009; Nishioka et al., 2008; Yagi et al., 2007). Among the four Tead members, Tead1 and Tead4 are highly expressed in cardiac and skeletal muscle (Jacquemin et al., 1996; Joshi et al., 2017; Wang et al., 2018). Tead1-null mice are embryonic lethal due to defects in cardiac remodeling and fetal heart development (Butler and Ordahl, 1999; Chen et al., 1994; Tan et al., 2020). Tead1 is also required for maintaining adult cardiomyocyte function, and its loss results in lethal dilated cardiomyopathy in mice and heart failure in humans (Liu et al., 2017; Tan et al., 2020). Tead4 is expressed in developing skeletal muscle in mouse embryos, and it is a direct target of MYOD1 and MYOG transcription factors in the C2C12 mouse myoblast cell line (Blais et al., 2005; Jacquemin et al., 1996). Further, cardiac muscle-specific transgenic overexpression of Tead4 has been shown to induce cardiac contractile dysfunction (Chen et al., 2004). In addition, many studies have reported crosstalk between Tead4 and non-coding RNAs in various cancers and muscle development, suggesting its critical role in different biological contexts (Chen et al., 2020b; Li et al., 2016; Tan et al., 2019). We found that *Platr4* directly interacts with Tead4 and our ChIP-qPCR result indicated a high occupancy of Tead4 on the *Ctgf* promoter, supporting its functional role in partnering with *Platr4* to regulate the *Ctgf* gene. Further, both gain-of-function and loss-of-function of Tead4, resulted in altered expression (upregulation or downregulation, respectively) of Ctgf, indicating a transcriptional regulatory role of Tead4 in this context. Thus, our results for the first time showed that the interaction of Tead4 with *Platr4* is essential for transcriptional regulation of Ctgf, thereby impacting the cardiac lineage. Future studies will examine the potential role of *Plat4* regulating other gene targets as well as the regulation of Tead4 gene expression programs.

Cardiovascular disease is the leading cause of death worldwide, although the adult mammalian heart has a finite ability in proliferation and regeneration (Bergmann et al., 2009). Therefore, understanding the molecular pathways of cardiogenesis, especially cardiac lineage differentiation, is crucial to study regenerative medicine. Stem cell models hold a great promise to look at this disease in a lab-dish and a potential for regenerative therapies (Zakrzewski et al., 2019). Based upon synteny, *Platr4* has three potential human orthologues RP11-130C6.1, RP11-184M15.1 and RP11-184M15.2, which will be the subject of future study. They are located in chromosome 4 (q28.1-28.3). Clinical case studies reported that deletion of chromosome 4 (q28.1-31.3) is associated with cardiac defects including arrhythmia, ventricular or atrial septal defects, thickening (hypertrophy) of the heart muscle (myocardium), and structural heart defects (Copelli et al., 1995; Frappaz et al., 1983; Ockey et al., 1967). Thus, we anticipate that one or more of these potential human lncRNA transcripts may be the human ortholog of *Platr4* and ideal candidates to explore their functional role in heart development and cardiac function.

## Author contributions

RH designed experiments, methodology, analyze data, investigation, and wrote the manuscript. LB methodology and editing. BB, NC methodology. LG, SL methodology, analyze data. MS writing - review and editing. JW histology analysis. DS conceptualization, supervision, funding acquisition, writing - review, and editing.

## Acknowledgments

We thank all members of the Spector laboratory for important suggestions throughout the course of this research. We would also like to thank the CSHL Shared Resources (Microscopy, Mass Spectrometry, Flow Cytometry, Animal, Histology, and Next-Gen Sequencing) for services and technical expertise (NCI 2P3OCA45508). This research was supported by NIGMS MIRA 65000201 (DLS) and by NIH R01 HD085904 (MMS).

## Declaration of interests

The authors declare no competing interests.

## Notes

### Competing Interest Statement

The authors have declared no competing interest.

## References

Aban, C.E., Lombardi, A., Neiman, G., Biani, M.C., La Greca, A., Waisman, A., Moro, L.N., Sevlever, G., Miriuka, S., and Luzzani, C. (2021). Downregulation of E-cadherin in pluripotent stem cells triggers partial EMT. Sci Rep 11, 2048.

Akizawa, H., Yanagawa, Y., Nagano, M., Bai, H., Takahashi, M., and Kawahara, M. (2019). Significance of CCN2 expression in bovine preimplantation development. Anim Sci J 90, 49–54.

Alexanian, M., Maric, D., Jenkinson, S.P., Mina, M., Friedman, C.E., Ting, C.C., Micheletti, R., Plaisance, I., Nemir, M., Maison, D., et al. (2017). A transcribed enhancer dictates mesendoderm specification in pluripotency. Nat Commun 8, 1806.

Anbanandam, A., Albarado, D.C., Nguyen, C.T., Halder, G., Gao, X., and Veeraraghavan, S. (2006). Insights into transcription enhancer factor 1 (TEF-1) activity from the solution structure of the TEA domain. Proc Natl Acad Sci U S A 103, 17225–17230.

Ballarino, M., Cipriano, A., Tita, R., Santini, T., Desideri, F., Morlando, M., Colantoni, A., Carrieri, C., Nicoletti, C., Musaro, A., et al. (2018). Deficiency in the nuclear long noncoding RNA Charme causes myogenic defects and heart remodeling in mice. EMBO J 37.

Benhaddou, A., Keime, C., Ye, T., Morlon, A., Michel, I., Jost, B., Mengus, G., and Davidson, I. (2012). Transcription factor TEAD4 regulates expression of myogenin and the unfolded protein response genes during C2C12 cell differentiation. Cell Death Differ 19, 220–231.

Bergmann, J.H., Li, J., Eckersley-Maslin, M.A., Rigo, F., Freier, S.M., and Spector, D.L. (2015). Regulation of the ESC transcriptome by nuclear long noncoding RNAs. Genome Res 25, 1336–1346.

Bergmann, O., Bhardwaj, R.D., Bernard, S., Zdunek, S., Barnabe-Heider, F., Walsh, S., Zupicich, J., Alkass, K., Buchholz, B.A., Druid, H., et al. (2009). Evidence for cardiomyocyte renewal in humans. Science 324, 98–102.

Blais, A., Tsikitis, M., Acosta-Alvear, D., Sharan, R., Kluger, Y., and Dynlacht, B.D. (2005). An initial blueprint for myogenic differentiation. Genes Dev 19, 553–569.

Butler, A.J., and Ordahl, C.P. (1999). Poly(ADP-ribose) polymerase binds with transcription enhancer factor 1 to MCAT1 elements to regulate muscle-specific transcription. Mol Cell Biol 19, 296–306.

Chatzizacharias, N.A., Kouraklis, G.P., and Theocharis, S.E. (2010). The role of focal adhesion kinase in early development. Histol Histopathol 25, 1039–1055.

Chen, H.H., Baty, C.J., Maeda, T., Brooks, S., Baker, L.C., Ueyama, T., Gursoy, E., Saba, S., Salama, G., London, B., et al. (2004). Transcription enhancer factor-1-related factor-transgenic mice develop cardiac conduction defects associated with altered connexin phosphorylation. Circulation 110, 2980–2987.

Chen, J., Wang, Y., Wang, C., Hu, J.F., and Li, W. (2020a). LncRNA Functions as a New Emerging Epigenetic Factor in Determining the Fate of Stem Cells. Front Genet 11, 277.

Chen, M., Huang, B., Zhu, L., Chen, K., Liu, M., and Zhong, C. (2020b). Structural and Functional Overview of TEAD4 in Cancer Biology. Onco Targets Ther 13, 9865–9874.

Chen, M.M., Lam, A., Abraham, J.A., Schreiner, G.F., and Joly, A.H. (2000). CTGF expression is induced by TGF-beta in cardiac fibroblasts and cardiac myocytes: a potential role in heart fibrosis. J Mol Cell Cardiol 32, 1805–1819.

Chen, Z., Friedrich, G.A., and Soriano, P. (1994). Transcriptional enhancer factor 1 disruption by a retroviral gene trap leads to heart defects and embryonic lethality in mice. Genes Dev 8, 2293–2301.

Copelli, S., del Rey, G., Heinrich, J., and Coco, R. (1995). Brief clinical report: interstitial deletion of the long arm of chromosome 4, del(4)(q28-->q31.3). Am J Med Genet 55, 77–79.

Croci, S., Landuzzi, L., Astolfi, A., Nicoletti, G., Rosolen, A., Sartori, F., Follo, M.Y., Oliver, N., De Giovanni, C., Nanni, P., et al. (2004). Inhibition of connective tissue growth factor (CTGF/CCN2) expression decreases the survival and myogenic differentiation of human rhabdomyosarcoma cells. Cancer Res 64, 1730–1736.

Daniels, A., van Bilsen, M., Goldschmeding, R., van der Vusse, G.J., and van Nieuwenhoven, F.A. (2009). Connective tissue growth factor and cardiac fibrosis. Acta Physiol (Oxf) 195, 321–338.

Derrien, T., Johnson, R., Bussotti, G., Tanzer, A., Djebali, S., Tilgner, H., Guernec, G., Martin, D., Merkel, A., Knowles, D.G., et al. (2012). The GENCODE v7 catalog of human long noncoding RNAs: analysis of their gene structure, evolution, and expression. Genome Res 22, 1775–1789.

Desbaillets, I., Ziegler, U., Groscurth, P., and Gassmann, M. (2000). Embryoid bodies: an in vitro model of mouse embryogenesis. Exp Physiol 85, 645–651.

Fagnocchi, L., Cherubini, A., Hatsuda, H., Fasciani, A., Mazzoleni, S., Poli, V., Berno, V., Rossi, R.L., Reinbold, R., Endele, M., et al. (2016). A Myc-driven self-reinforcing regulatory network maintains mouse embryonic stem cell identity. Nat Commun 7, 11903.

Frappaz, D., Bourgeois, J., Berthier, J.C., Laurent, C., and Bethenod, M. (1983). [Syndrome of terminal deletion of the long arm of chromosome 4. Apropos of a personal case with a review of the literature]. Pediatrie 38, 261–270.

Gardin, J.M., Siri, F.M., Kitsis, R.N., Edwards, J.G., and Leinwand, L.A. (1995). Echocardiographic assessment of left ventricular mass and systolic function in mice. Circ Res 76, 907–914.

Gerritsen, K.G., Falke, L.L., van Vuuren, S.H., Leeuwis, J.W., Broekhuizen, R., Nguyen, T.Q., de Borst, G.J., Nathoe, H.M., Verhaar, M.C., Kok, R.J., et al. (2016). Plasma CTGF is independently related to an increased risk of cardiovascular events and mortality in patients with atherosclerotic disease: the SMART study. Growth Factors 34, 149–158.

Goh, S.K., Olsen, P., and Banerjee, I. (2013). Extracellular matrix aggregates from differentiating embryoid bodies as a scaffold to support ESC proliferation and differentiation. PLoS One 8, e61856.

Grote, P., Wittler, L., Hendrix, D., Koch, F., Wahrisch, S., Beisaw, A., Macura, K., Blass, G., Kellis, M., Werber, M., et al. (2013). The tissue-specific lncRNA Fendrr is an essential regulator of heart and body wall development in the mouse. Dev Cell 24, 206–214.

Guo, X., Xu, Y., Wang, Z., Wu, Y., Chen, J., Wang, G., Lu, C., Jia, W., Xi, J., Zhu, S., et al. (2018). A Linc1405/Eomes Complex Promotes Cardiac Mesoderm Specification and Cardiogenesis. Cell Stem Cell 22, 893–908 e896.

Gustafsson, E., and Fassler, R. (2000). Insights into extracellular matrix functions from mutant mouse models. Exp Cell Res 261, 52–68.

Guttman, M., Donaghey, J., Carey, B.W., Garber, M., Grenier, J.K., Munson, G., Young, G., Lucas, A.B., Ach, R., Bruhn, L., et al. (2011). lincRNAs act in the circuitry controlling pluripotency and differentiation. Nature 477, 295–300.

Han, P., Li, W., Lin, C.H., Yang, J., Shang, C., Nuernberg, S.T., Jin, K.K., Xu, W., Lin, C.Y., Lin, C.J., et al. (2014). A long noncoding RNA protects the heart from pathological hypertrophy. Nature 514, 102–106.

Han, X., Luo, S., Peng, G., Lu, J.Y., Cui, G., Liu, L., Yan, P., Yin, Y., Liu, W., Wang, R., et al. (2018). Mouse knockout models reveal largely dispensable but context-dependent functions of lncRNAs during development. J Mol Cell Biol 10, 175–178.

Hobuss, L., Bar, C., and Thum, T. (2019). Long Non-coding RNAs: At the Heart of Cardiac Dysfunctionã Front Physiol 10, 30.

Hoffman, M., Kyriazis, I.D., Lucchese, A.M., de Lucia, C., Piedepalumbo, M., Bauer, M., Schulze, P.C., Bonios, M.J., Koch, W.J., and Drosatos, K. (2019). Myocardial Strain and Cardiac Output are Preferable Measurements for Cardiac Dysfunction and Can Predict Mortality in Septic Mice. J Am Heart Assoc 8, e012260.

Hou, N., Wen, Y., Yuan, X., Xu, H., Wang, X., Li, F., and Ye, B. (2017). Activation of Yap1/Taz signaling in ischemic heart disease and dilated cardiomyopathy. Exp Mol Pathol 103, 267–275.

Jacquemin, P., Hwang, J.J., Martial, J.A., Dolle, P., and Davidson, I. (1996). A novel family of developmentally regulated mammalian transcription factors containing the TEA/ATTS DNA binding domain. J Biol Chem 271, 21775–21785.

Jacquemin, P., Martial, J.A., and Davidson, I. (1997). Human TEF-5 is preferentially expressed in placenta and binds to multiple functional elements of the human chorionic somatomammotropin-B gene enhancer. J Biol Chem 272, 12928–12937.

Janky, R., Verfaillie, A., Imrichova, H., Van de Sande, B., Standaert, L., Christiaens, V., Hulselmans, G., Herten, K., Naval Sanchez, M., Potier, D., et al. (2014). iRegulon: from a gene list to a gene regulatory network using large motif and track collections. PLoS Comput Biol 10, e1003731.

Joshi, S., Davidson, G., Le Gras, S., Watanabe, S., Braun, T., Mengus, G., and Davidson, I. (2017). TEAD transcription factors are required for normal primary myoblast differentiation in vitro and muscle regeneration in vivo. PLoS Genet 13, e1006600.

Kang, W., Huang, T., Zhou, Y., Zhang, J., Lung, R.W.M., Tong, J.H.M., Chan, A.W.H., Zhang, B., Wong, C.C., Wu, F., et al. (2018). miR-375 is involved in Hippo pathway by targeting YAP1/TEAD4-CTGF axis in gastric carcinogenesis. Cell Death Dis 9, 92.

Kim, J.M., Moon, S.H., Lee, S.G., Cho, Y.J., Hong, K.S., Lee, J.H., Lee, H.J., and Chung, H.M. (2011). Assessment of differentiation aspects by the morphological classification of embryoid bodies derived from human embryonic stem cells. Stem Cells Dev 20, 1925–1935.

Klattenhoff, C.A., Scheuermann, J.C., Surface, L.E., Bradley, R.K., Fields, P.A., Steinhauser, M.L., Ding, H., Butty, V.L., Torrey, L., Haas, S., et al. (2013). Braveheart, a long noncoding RNA required for cardiovascular lineage commitment. Cell 152, 570–583.

Kleinman, H.K., Philp, D., and Hoffman, M.P. (2003). Role of the extracellular matrix in morphogenesis. Curr Opin Biotechnol 14, 526–532.

Koitabashi, N., Arai, M., Niwano, K., Watanabe, A., Endoh, M., Suguta, M., Yokoyama, T., Tada, H., Toyama, T., Adachi, H., et al. (2008). Plasma connective tissue growth factor is a novel potential biomarker of cardiac dysfunction in patients with chronic heart failure. Eur J Heart Fail 10, 373–379.

Kong, L., Zhang, Y., Ye, Z.Q., Liu, X.Q., Zhao, S.Q., Wei, L., and Gao, G. (2007). CPC: assess the protein-coding potential of transcripts using sequence features and support vector machine. Nucleic Acids Res 35, W345–349.

Lambi, A.G., Pankratz, T.L., Mundy, C., Gannon, M., Barbe, M.F., Richtsmeier, J.T., and Popoff, S.N. (2012). The skeletal site-specific role of connective tissue growth factor in prenatal osteogenesis. Dev Dyn 241, 1944–1959.

Lee, S.W., Won, J.Y., Kim, W.J., Lee, J., Kim, K.H., Youn, S.W., Kim, J.Y., Lee, E.J., Kim, Y.J., Kim, K.W., et al. (2013). Snail as a potential target molecule in cardiac fibrosis: paracrine action of endothelial cells on fibroblasts through snail and CTGF axis. Mol Ther 21, 1767–1777.

Leeuwis, J.W., Nguyen, T.Q., Theunissen, M.G., Peeters, W., Goldschmeding, R., Pasterkamp, G., and Vink, A. (2010). Connective tissue growth factor is associated with a stable atherosclerotic plaque phenotype and is involved in plaque stabilization after stroke. Stroke 41, 2979–2981.

Li, Z., Ouyang, H., Zheng, M., Cai, B., Han, P., Abdalla, B.A., Nie, Q., and Zhang, X. (2016). Integrated Analysis of Long Non-coding RNAs (LncRNAs) and mRNA Expression Profiles Reveals the Potential Role of LncRNAs in Skeletal Muscle Development of the Chicken. Front Physiol 7, 687.

Lim, J., and Thiery, J.P. (2012). Epithelial-mesenchymal transitions: insights from development. Development 139, 3471–3486.

Lin, M.F., Jungreis, I., and Kellis, M. (2011). PhyloCSF: a comparative genomics method to distinguish protein coding and non-coding regions. Bioinformatics 27, i275–282.

Liu, R., Lee, J., Kim, B.S., Wang, Q., Buxton, S.K., Balasubramanyam, N., Kim, J.J., Dong, J., Zhang, A., Li, S., et al. (2017). Tead1 is required for maintaining adult cardiomyocyte function, and its loss results in lethal dilated cardiomyopathy. JCI Insight 2.

Liu, X., Li, H., Rajurkar, M., Li, Q., Cotton, J.L., Ou, J., Zhu, L.J., Goel, H.L., Mercurio, A.M., Park, J.S., et al. (2016a). Tead and AP1 Coordinate Transcription and Motility. Cell Rep 14, 1169–1180.

Liu, Y., Chen, L., Diaz, A.D., Benham, A., Xu, X., Wijaya, C.S., Fa’ak, F., Luo, W., Soibam, B., Azares, A., et al. (2016b). Mesp1 Marked Cardiac Progenitor Cells Repair Infarcted Mouse Hearts. Sci Rep 6, 31457.

Lockhart, M., Wirrig, E., Phelps, A., and Wessels, A. (2011). Extracellular matrix and heart development. Birth Defects Res A Clin Mol Teratol 91, 535–550.

Luo, Q., Kang, Q., Si, W., Jiang, W., Park, J.K., Peng, Y., Li, X., Luu, H.H., Luo, J., Montag, A.G., et al. (2004). Connective tissue growth factor (CTGF) is regulated by Wnt and bone morphogenetic proteins signaling in osteoblast differentiation of mesenchymal stem cells. J Biol Chem 279, 55958–55968.

Mercer, T.R., Qureshi, I.A., Gokhan, S., Dinger, M.E., Li, G., Mattick, J.S., and Mehler, M.F. (2010). Long noncoding RNAs in neuronal-glial fate specification and oligodendrocyte lineage maturation. BMC Neurosci 11, 14.

Morrison, B.L., Jose, C.C., and Cutler, M.L. (2010). Connective Tissue Growth Factor (CTGF/CCN2) enhances lactogenic differentiation of mammary epithelial cells via integrin-mediated cell adhesion. BMC Cell Biol 11, 35.

Murry, C.E., and Keller, G. (2008). Differentiation of embryonic stem cells to clinically relevant populations: lessons from embryonic development. Cell 132, 661–680.

Nishioka, N., Inoue, K., Adachi, K., Kiyonari, H., Ota, M., Ralston, A., Yabuta, N., Hirahara, S., Stephenson, R.O., Ogonuki, N., et al. (2009). The Hippo signaling pathway components Lats and Yap pattern Tead4 activity to distinguish mouse trophectoderm from inner cell mass. Dev Cell 16, 398–410.

Nishioka, N., Yamamoto, S., Kiyonari, H., Sato, H., Sawada, A., Ota, M., Nakao, K., and Sasaki, H. (2008). Tead4 is required for specification of trophectoderm in pre-implantation mouse embryos. Mech Dev 125, 270–283.

Ockey, C.H., Feldman, G.V., Macaulay, M.E., and Delaney, M.J. (1967). A large deletion of the long arm of chromosome No. 4 in a child with limb abnormalities. Arch Dis Child 42, 428–434.

Qi, X., Yang, G., Yang, L., Lan, Y., Weng, T., Wang, J., Wu, Z., Xu, J., Gao, X., and Yang, X. (2007). Essential role of Smad4 in maintaining cardiomyocyte proliferation during murine embryonic heart development. Dev Biol 311, 136–146.

Ramazani, Y., Knops, N., Elmonem, M.A., Nguyen, T.Q., Arcolino, F.O., van den Heuvel, L., Levtchenko, E., Kuypers, D., and Goldschmeding, R. (2018). Connective tissue growth factor (CTGF) from basics to clinics. Matrix Biol 68-69, 44–66.

Ramos, A.D., Andersen, R.E., Liu, S.J., Nowakowski, T.J., Hong, S.J., Gertz, C., Salinas, R.D., Zarabi, H., Kriegstein, A.R., and Lim, D.A. (2015). The long noncoding RNA Pnky regulates neuronal differentiation of embryonic and postnatal neural stem cells. Cell Stem Cell 16, 439–447.

Rickard, A.J., Morgan, J., Tesch, G., Funder, J.W., Fuller, P.J., and Young, M.J. (2009). Deletion of mineralocorticoid receptors from macrophages protects against deoxycorticosterone/salt-induced cardiac fibrosis and increased blood pressure. Hypertension 54, 537–543.

Riley, J.K., Carayannopoulos, M.O., Wyman, A.H., Chi, M., Ratajczak, C.K., and Moley, K.H. (2005). The PI3K/Akt pathway is present and functional in the preimplantation mouse embryo. Dev Biol 284, 377–386.

Rinn, J.L., and Chang, H.Y. (2012). Genome regulation by long noncoding RNAs. Annu Rev Biochem 81, 145–166.

Rozario, T., and DeSimone, D.W. (2010). The extracellular matrix in development and morphogenesis: a dynamic view. Dev Biol 341, 126–140.

Sart, S., Ma, T., and Li, Y. (2014). Extracellular matrices decellularized from embryonic stem cells maintained their structure and signaling specificity. Tissue Eng Part A 20, 54–66.

Shi-Wen, X., Leask, A., and Abraham, D. (2008). Regulation and function of connective tissue growth factor/CCN2 in tissue repair, scarring and fibrosis. Cytokine Growth Factor Rev 19, 133–144.

Slaymaker, I.M., Gao, L., Zetsche, B., Scott, D.A., Yan, W.X., and Zhang, F. (2016). Rationally engineered Cas9 nucleases with improved specificity. Science 351, 84–88.

Smith, K.N., Miller, S.C., Varani, G., Calabrese, J.M., and Magnuson, T. (2019). Multimodal Long Noncoding RNA Interaction Networks: Control Panels for Cell Fate Specification. Genetics 213, 1093–1110.

Song, L., Yan, W., Chen, X., Deng, C.X., Wang, Q., and Jiao, K. (2007). Myocardial smad4 is essential for cardiogenesis in mouse embryos. Circ Res 101, 277–285.

Tan, B.S., Yang, M.C., Singh, S., Chou, Y.C., Chen, H.Y., Wang, M.Y., Wang, Y.C., and Chen, R.H. (2019). LncRNA NORAD is repressed by the YAP pathway and suppresses lung and breast cancer metastasis by sequestering S100P. Oncogene 38, 5612–5626.

Tan, W.L.W., Anene-Nzelu, C.G., Wong, E., Lee, C.J.M., Tan, H.S., Tang, S.J., Perrin, A., Wu, K.X., Zheng, W., Ashburn, R.J., et al. (2020). Epigenomes of Human Hearts Reveal New Genetic Variants Relevant for Cardiac Disease and Phenotype. Circ Res 127, 761–777.

Tsika, R.W., Ma, L., Kehat, I., Schramm, C., Simmer, G., Morgan, B., Fine, D.M., Hanft, L.M., McDonald, K.S., Molkentin, J.D., et al. (2010). TEAD-1 overexpression in the mouse heart promotes an age-dependent heart dysfunction. J Biol Chem 285, 13721–13735.

von Gise, A., Lin, Z., Schlegelmilch, K., Honor, L.B., Pan, G.M., Buck, J.N., Ma, Q., Ishiwata, T., Zhou, B., Camargo, F.D., et al. (2012). YAP1, the nuclear target of Hippo signaling, stimulates heart growth through cardiomyocyte proliferation but not hypertrophy. Proc Natl Acad Sci U S A 109, 2394–2399.

Wang, J., Zhang, F., Yang, H., Wu, H., Cui, R., Zhao, Y., Jiao, C., Wang, X., Liu, X., Wu, L., et al. (2018). Effect of TEAD4 on multilineage differentiation of muscle-derived stem cells. Am J Transl Res 10, 998–1011.

Wang, K.C., and Chang, H.Y. (2011). Molecular mechanisms of long noncoding RNAs. Mol Cell 43, 904–914.

Wang, L., Park, H.J., Dasari, S., Wang, S., Kocher, J.P., and Li, W. (2013). CPAT: Coding-Potential Assessment Tool using an alignment-free logistic regression model. Nucleic Acids Res 41, e74.

Xin, M., Kim, Y., Sutherland, L.B., Qi, X., McAnally, J., Schwartz, R.J., Richardson, J.A., Bassel-Duby, R., and Olson, E.N. (2011). Regulation of insulin-like growth factor signaling by Yap governs cardiomyocyte proliferation and embryonic heart size. Sci Signal 4, ra70.

Yagi, R., Kohn, M.J., Karavanova, I., Kaneko, K.J., Vullhorst, D., DePamphilis, M.L., and Buonanno, A. (2007). Transcription factor TEAD4 specifies the trophectoderm lineage at the beginning of mammalian development. Development 134, 3827–3836.

Yang, J., Mani, S.A., Donaher, J.L., Ramaswamy, S., Itzykson, R.A., Come, C., Savagner, P., Gitelman, I., Richardson, A., and Weinberg, R.A. (2004). Twist, a master regulator of morphogenesis, plays an essential role in tumor metastasis. Cell 117, 927–939.

Zakrzewski, W., Dobrzynski, M., Szymonowicz, M., and Rybak, Z. (2019). Stem cells: past, present, and future. Stem Cell Res Ther 10, 68.

Zhang, J., Klos, M., Wilson, G.F., Herman, A.M., Lian, X., Raval, K.K., Barron, M.R., Hou, L., Soerens, A.G., Yu, J., et al. (2012). Extracellular matrix promotes highly efficient cardiac differentiation of human pluripotent stem cells: the matrix sandwich method. Circ Res 111, 1125–1136.

Zhao, B., Ye, X., Yu, J., Li, L., Li, W., Li, S., Yu, J., Lin, J.D., Wang, C.Y., Chinnaiyan, A.M., et al. (2008). TEAD mediates YAP-dependent gene induction and growth control. Genes Dev 22, 1962–1971.

Zhou, Y., Huang, T., Cheng, A.S., Yu, J., Kang, W., and To, K.F. (2016). The TEAD Family and Its Oncogenic Role in Promoting Tumorigenesis. Int J Mol Sci 17.

