## supplements for "*Platr4* is an ESC-specific lncRNA that exhibits its function downstream on meso/endoderm lineage commitment"

### Extended Experimental Methods

**ESC lines:**
Mouse embryonic stem cells (ESC) AB2.2 (Gift from Dr. Alea Mills) and V6.5 (CSHL Gene Targeting Facility) were maintained on an irradiated feeder layer (Millipore) in knockout DMEM medium supplemented with 15% FBS, sodium pyruvate, non-essential amino acids, β-mercaptoethanol and leukemia inhibiting factor (Millipore). All cell culture reagents were obtained from Gibco (Life Technologies) unless otherwise stated.

**ESC differentiation into EBs:**
Feeder cells were depleted from ESCs by one-hour soaking and resuspended to 20,000 cells/ml in a complete growth medium without leukaemia inhibitory factor (LIF). The suspension was plated as 20ul drops on the lid of the Petri dish, which were then inverted and incubated for two days to induce aggregation by the hanging drop method. Aggregated ESCs formed the embryoid body (EB). EBs were then collected and plated on standard gelatin-coated 6-well cell culture plates in a complete growth medium without LIF for 12 days to analyze the percentage of spontaneous beating EBs. Further, EBs were further collected at different time points (day 0, 3, 5, 7, 9 and 12) to interpret gene expression and for H&E-staining. To differentiate into neuroectoderm lineage, EBs were grown in a complete growth media containing 1uM all-trans retinoic acid lacking LIF to differentiate them into the neuroectoderm lineage.

**Cardiomyocyte Differentiation:**
ESCs were directed to differentiate into cardiomyocytes as described previously (Klattenhoff et al., 2013). Briefly, cells were depleted from the feeder layer with a standard technique and aggregated into EBs using the hanging drop method. Next, EBs were dissociated and cultured at a density of 100,000 cells/ml for two days in serum-free media (3 parts IMDM (Cellgro): 1 part Ham's F12 (Gibco), 0.05% BSA, 2 mM GlutaMax (Gibco), B27 supplement (Gibco), N2 supplement (Gibco) supplemented with 50 mg/ml ascorbic acid and 4.5 x$10^{-4}$ M monothioglycerol (Sigma). Around 48 hours later, EBs were dissociated and re-aggregated in the presence of 5 ng/mL hVEGF (R&D) and hActivin A (5 ng/ml) (R&D) and hBMP4 (0.25 ng/ml) (R&D). EBs were further dissociated and replated at 500,000 cells/well of 24 well plate in StemPro-34 (Gibco) supplemented with 5 ng/mL hVEGF, 10 ng/mL human basic FGF (R&D) and 25 ng/mL FGF10 (R&D).

**CRISPR/Cas9-deleted ESC lines:**
Two guide RNAs (sgRNAs) targeting the transcription start site (TSS) of Platr4 were placed into the pSpCas9(BB)-2A-GFP (PX458) vector (Addgene # 48138). The sgRNAs were designed using http://crispr.mit.edu/. ESCs were transfected with plasmids using 4D-Nucleofector X Unit, program code CG104. Transfected ESCs were sorted 48 hours post-transfection, as single-cell deposition into 96-well plates using a FACS Aria (SORP) Cell Sorter (BD). Each cell clone was cultured and examined by genomic PCR, qRT-PCR, and Sanger sequencing to select a homozygous knockout clone. Cells were transfected with a sgRNA targeting Renilla luciferase, used as a negative control. sgRNAs sequence and primers are provided in Supplementary data 1.

**5’/3' Rapid Amplification of cDNA Ends (RACE):**
5′ and 3′ RACE was performed using the Ambion FirstChoice RLM-RACE kit according to the manufacturer's protocol. Briefly, fragments were amplified by nested PCR using AmpliTaq Polymerase, and PCR products were cloned into pGEM-T Easy kit (Promega), and clones were sequenced using standard Sanger sequencing. See Supplementary data 1 for primer sequences.

**Northern blot Analysis:**
5 µg of polyA+ enriched RNA (Dynabeads mRNA purification kit, Invitrogen) was resolved on a 1% denaturing polyacrylamide gel and transferred to a Hybond XL membrane (GE Healthcare Life Sciences) and crosslinked (Stratalinker 1800 UV, Stratagene). The membrane was prehybridized with ULTRAhyb buffer (Ambion) for two hours, then hybridized with the Platr4 specific radiolabeled DNA probe overnight at $42^0$C. Platr4-specific DNA probes were labeled with [$\gamma$-$^{32}$P] ATP in a random primed labeling reaction using the Prime-It RmT Random Primer Labeling Kit (Stratagene). The next day hybridized blot was washed three times in 2xSSC/0.1%SDS and once in 0.1xSSC/0.1%SDS. Signal was quantified using the Fujifilm Life Science FLA-5100 imaging system and X-ray film.

**Western blot analysis:**
Cells were washed with PBS and lysed in RIPA buffer (25 mM Tris-HCl pH 7.6, 150 mM NaCl, 1% NP-40 substitute, 1% sodium deoxycholate, and 0.1% SDS) supplemented with 1X Roche protease inhibitor cocktail on ice for 30 min with occasional vortex. Lysate was then centrifuged at 13000xg for 30 min. The supernatant was collected, and protein concentration was measured by BCA protein assay. Proteins were separated by SDS-PAGE on 4%–20% discontinuous polyacrylamide gels. The proteins were transferred onto a nitrocellulose membrane. The membrane was blocked in 5% non-fat dry milk for 1 hour at room temperature and incubated with primary antibodies (see below) at 4°C overnight. The following day membranes were washed three times for 5 mins each in PBS-Tween (0.05%) and incubated with appropriate secondary antibody for 1 hour at room temperature. Then washed the membrane three times and the signal was visualized using west pico chemiluminescent substrate (Thermo Scientific). The primary antibodies are as follows: anti-Tead4 (Abcam, ab97460), anti-Tead1 (BD Biosciences, 610923), anti-Tead2 (Biorbyt, orb382464), anti-Tead3 (Novus Biologicals, NBP1-83949), anti-Ctgf (Novus Biologicals, NB100-724SS), anti-Actin (Sigma, A5441) was used as the loading control.

**RNA Isolation and Quantitative Real-Time PCR (qRT-PCR) Assays:**
Total RNA was extracted from cells or tissues using TRIzol according to the manufacturer's instruction. 1 µg total RNA was used to synthesize cDNA using the TaqMan Reverse Transcription Reagent kit. 30ng of cDNA was used to perform qRT-PCR reaction using SYBR green PCR master mix on an ABI QuantStudio 6 Flex Real-Time PCR System. The housekeeping genes Gapdh, CycloB, and Pabpc were used as internal controls to normalize the gene of interest. All the experiments were performed in duplicates and repeated three times. Primers sequences were listed in Supplementary data 1.

**Cell Fractionation:**
Cell fractionation was performed as previously described (Conrad and Orom, 2017). Briefly, 5 million cells were resuspended in ice-cold lysis buffer containing 10 mM Tris pH 7.5, 150 mM NaCl and 0.15% NP-40 substitute. The cytoplasmic fraction was separated from nuclei by overlaid

the cell suspension on the sucrose buffer containing 10 mM Tris pH7.4, 150 mM NaCl, and 24% sucrose and centrifuged at 3500xg for 10 minutes. The remaining nuclei pellet was rinsed with ice-cold PBS-EDTA once and resuspended in urea buffer (1 M Urea, 0.3 M NaCl, 7.5 mM $MgCl_2$, 0.2 mM EDTA, and 1% NP-40 substitute) on ice for 2 minutes. The lysate was then centrifuged at 13000xg for 2 minutes to separate the chromatin pellet from the supernatant comprising the nucleoplasm fraction. The cytoplasm fraction, nucleoplasm fraction, and chromatin pellet were used for RNA extraction using TRIzol reagent according to the manufacturer's protocol.

**Cloning:**
For overexpression of specific genes, an insert containing the gene of interest was cloned into pBApo-EF1alpha Puro (TaKaRa) and pCMV6-entry (Origene) plasmids using the manufacturer's protocol. Plasmids were transformed into NEB Stable competent *E. coli* using the heat shock method. Four to five colonies per plate were picked and sequenced using standard Sanger sequencing.

**Knockdown (KD) using siRNA:**
The siRNA transfection was performed in ESCs using 4D-Nucleofector X Unit, program code CG104. ON-TARGET-plus SMARTpool siRNAs for Tead1, Tead2, Tead4 and non-targeting siRNA were purchased from Dharmacon Inc. (Chicago, Il., USA). The experiments were performed at least in triplicates.

**RNA sequencing and analysis:**
Total RNA was extracted from both ESCs and EBs using TRIzol according to the manufacturer's instruction. RNA quality was assayed by running an RNA 6000 Nanochip on a 2100 Bioanalyzer. Each RNA sample had an RNA integrity number (RIN) of 9 or above. 500ng total RNA was used to prepare poly(A)+ enriched RNA-seq libraries using the Illumina TruSeq sample prep kit following the manufacture's protocol. The libraries were multiplexed and sequenced single-end 75 bp on the NextSeq500 platform (Illumina). Reads were then mapped to the mouse mm10 genome using RNA STAR (Dobin et al., 2013), and reads per gene record were counted using HTSeq-counts (Andres et al., 2015)) using the GENCODE M20 annotation. The list of differentially expressed genes was generated using DESeq2 (Love et al., 2014), and an FDR-adjusted P-value of <0.05 was set as a threshold for statistical significance. The KEGG pathway and GO term enrichment were carried out using the R/Bioconductor packages GAGE (Luo et al., 2009) and Pathview (Luo and Brouwer, 2013).

**iRegulon analysis:**
iRegulon v1.3 (build: 2015-02-12), analysis was performed as described previously (Janky et al., 2014). The master regulator transcription factors were identified using a differential gene set as input for motif search. The thresholds set for this motif enrichment analysis were as follows: the minimum normalized enrichment score (NES) > 3, false discovery rate (FDR) on motif similarity < 0.001. Motif collection was set to 10 K (9713 PWMs), and putative regulatory search regions of 10kb centered around the TSS (7 species) and 500bp upstream of TSS for the ranking option. Targeted transcription factors were obtained based on HOMER, TRANSFAC, yeTFaSCo databases included in iRegulon analysis.

**RNA immunoprecipitation (RIP):**
RIP was performed as previously described (Zhang et al., 2016). Briefly, 10 million ESCs were harvested in tryple E, washed with cold PBS, resuspended in 2 ml PBS, 2 ml nuclear isolation buffer (1.28 M sucrose, 40 mM Tris-HCl pH 7.5, 20 mM MgCl2, 4% Triton X-100), and 6 ml nuclease-free water. Incubate the cells on ice for 20 min with intermittent vortex. The lysate was pelleted by centrifugation at 2500×g for 15 min to harvest nuclei. Nuclei pellets were resuspended in 1 ml RIP buffer (150 mM KCl, 25 mM Tris pH 7.5, 5 mM EDTA, 0.5 mM DTT, 0.5% NP-40, 100 U/ml SUPERase-IN, and 1X Roche protease inhibitor cocktail) and sonicated for 15 min using Pico BioRuptor (30 s ON/OFF) (Diagenode) at 4 °C. The lysate was then centrifuged at 16,000×g for 10 min. The supernatant was collected and incubated with 4 μg Tead4 (Abcam # ab58310), Tead1 (Cell signaling # 12292), Tead2 (Biorbyt # orb382464), Tead3 (Novus Biologicals # NBP1-8394) antibodies mouse or rabbit isotype IgG control overnight at 4 °C with gentle shaking. The next day, 40 μl of protein G and A beads (Thermo Fisher) for mouse and rabbit antibodies respectively were added into the immune-complex reactions and incubated for two hours at 4 °C with gentle shaking. Beads were washed three times with RIP buffer and once with PBS. Then beads were collected for western blot analysis and RNA extraction to perform qRT-PCR—primer sequences listed in Supplementary data 1.

**Chromatin immunoprecipitation (ChIP) coupled with quantitative PCR (ChIP-qPCR):**
ChIP was performed according to the manufacturer's protocol (Active Motif) with minor modification. 15 million cells were crosslinked with 1% formaldehyde for 10 min at room temperature. The reaction was then quenched by adding glycine at a final concentration of 0.125M at room temperature for 5 min. Cells were centrifuged at 2500 rpm for 10 min at 4 °C, and the supernatant was discarded. The cell pellet was resuspended in lysis buffer and incubated on ice for 30 min followed by a centrifugation step at 5000 rpm for 10 mins at 4 °C to pellet the nuclei. Next, nuclei pellets were resuspended in shearing buffer and sonicated for 20 min using a Pico BioRuptor (30 s ON/OFF) at 4 °C. The sonicated sample was centrifuged at max speed for 15 min at 4 °C, and the supernatant was transferred into a new Eppendorf tube for IP. The antibodies used were Anti-Tead4 (Abcam # ab58310) was used for ChIP assay. ChIP-enriched DNA was quantified by qPCR on an ABI QuantStudio 6 Flex Real-Time PCR System. ChIP-qPCR primers can be found in Supplementary data 1.

**Embryo collection and in vitro culture:**
Embryos were collected at E0.5 day from swollen ampulas, treated with hyaluronidase (Sigma) at 37°C for 2–3 mins to remove cumulus cells. Embryos were then washed three times in PBS with BSA (Sigma, 6 mg/ml) and cultured for three days in 15μl drops of Potassium Simplex Optimized Medium (KSOM-AA, Millipore) covered with mineral oil (Ovoil, Vitrolife) in a humidified chamber at 37°C with 5% $CO_2$. Post-implantation embryos were collected at different embryonic days (E6.5, E8.5, E10.5, E12.5), fixed in 4% PFA overnight at 4°C, washed with PBS, and used for whole-mount RNA FISH.

**Single-molecule RNA fluorescence in situ hybridization (FISH):**

*ESCs:*
Single-molecule RNA-FISH was performed according to the manufacturer's protocol for the Affymetrix View ISH Cell Assay Kit (Thermo Fisher, QVT0050) using custom Type-6 primary

probes targeting Platr4. ESCs were seeded onto acid-cleaned #1.5 glass coverslips (Electron Microscopy Sciences, 72230-01) for 24 hours to reach 80% confluence, then fixed in freshly prepared 4% paraformaldehyde (PFA) (Electron Microscopy Sciences, 19200). Fixed cells were permeabilized and protease digested before hybridization. The hybridization and signal amplification steps were performed according to the manufacturer's instruction, and nuclei were counterstained with DAPI. Coverslips were mounted in ProLong Gold Antifade mounting medium (Thermo Fisher, P36930) for imaging on Zeiss LSM 710/780 Confocal Microscope systems.

*Whole-mount embryos:*
Single-molecule RNA FISH was performed on formalin-fixed whole-mount embryos using RNAscope® Fluorescent Multiplex Reagent Kit 320850 (ACD # 320850). Pretreatment of the tissue sections, hybridization, and signal amplification were all performed according to the manufacturer's instructions. Mounted embryos were imaged on a Zeiss LSM710 or LSM780 spinning disk confocal microscope.

**Immunofluorescence (IF):**

*ESCs:*
ESCs were cultured onto acid-cleaned #1.5 glass coverslips for 24 hours for 80-90% confluence. The following day cells were fixed in freshly prepared 4% paraformaldehyde (PFA) for 20 min at room temperature, washed three times with PBS, permeabilized in PBS with 0.2% Triton-X, 0.1% Tween-20 for 30 min on ice. Permeabilized cells were blocked in PBS with 1% BSA for one hour at room temperature and then incubated cells with anti-Oct4 antibody (mouse monoclonal, Santa Cruz, sc-5279, dilution 1:100) overnight at 4 °C. The next day, coverslips were washed three times in blocking reagent and stained with anti-mouse secondary (Alexafluor 488). After washing three-time, nuclei were counterstained with DAPI, and Coverslips were imaged on the Zeiss LSM 710/780 Confocal Microscope.

**Hematoxylin and Eosin (H&E) staining:**
Heart tissue and EBs were fixed in 4% PFA overnight, washed three times in PBS, and dehydrated by a series of ethanol and Xylene washes before embedding in Paraffin. Paraffin-embedded tissue and EBs were sectioned and stained with H&E. Slides were scanned using an Aperio slide scanner. A pathologist analyzed the sections to assess the histologic grade

**RNA Pull-Down:**
This experiment was performed as previously described (Klattenhoff et al., 2013). Briefly, 10 million cells were used to prepare nuclear pellets and the pellet was resuspended in RIP buffer (150 mM KCl, 25 mM Tris pH 7.5, 0.5 mM DTT, 0.5% NP40, 1mM ABSEF, protease inhibitor cocktail (Roche) and 20U/ml RNaseOut (Invitrogen)) and sonicated for 10 cycles using a Pico BioRuptor (30 s ON/OFF) at 4 °C. Sonicated lysate was then centrifuged for 10 min at 13,000 rpm. 50pmol of in vitro transcribed biotinylated RNA was added to the supernatant and incubated for one hour at room temperature, followed by addition of 60 µl streptavidin Dynabeads (Invitrogen) and incubated for one hour at room temperature. The lysate was then washed four times with RIP buffer, and beads were denatured in SDS buffer to release protein. Proteins were analyzed by Western blotting.

**Knockout mouse generation:**
All animal procedures and studies were approved by the Cold Spring Harbor Laboratory Animal Use Committee in accordance with IACUC procedures. Animals were maintained, and the experiments were performed with oversight of the Cold Spring Harbor Laboratory Animal Shared Resource, which is fully accredited by the Association for Assessment and Accreditation of Laboratory Animal Care (AAALAC). *Platr4* knockout mice were generated in the Gene Targeting and Transgenic Core facility at the University of Rochester Medical Centre (Han et al., 2014). Briefly, two sgRNAs (Supplementary data1) near the TSS were used to delete the promoter of Platr4 lncRNA. Guide RNA was in vitro transcribed by T7 promoter using MEGAshortscriptT7 Transcription Kit (Ambion, AM1354). In vitro transcribed RNAs were purified by MEGAclear Transcription Clean-Up Kit (Ambion, AM1908). Fertilized embryos were collected from oviducts of superovulated females. sgRNAs (50 ng/µl) and Cas9 mRNA (100 ng/µl) were co-injected into mouse zygotes with well-recognized pronuclei and then transferred into the uterus of pseudo-pregnant ICR females. Founder mice were tail-snipped for genotyping and sequencing. After confirming genomic deletion, each founder mice were successfully bred to the F1 generation using C57BL/6J mice for germline transmission. Heterozygous mice backcrossed to generate homozygous mice.

**Genotyping:**
Tail DNA was prepared using standard methods and analyzed by PCR. KO primer pairs were designed outside the deleted region, and WT primer pairs were created inside the deleted area (Supplemental data 1). KO PCR bands from each founder were confirmed by Sanger sequencing.

**Echocardiography:**
All cardiac ultrasound imaging was performed using a Vevo3100 scanner (VisualSonics) and the MX550D transducer. In preparation for the scan and on the same days as imaging, mice were anesthetized with 2-3% isoflurane. The hair over the chest region was entirely removed by shaving and Nair (depilation cream). For the actual scan, mice were again anesthetized with 2-3% isoflurane, and their paws lightly tapped onto the copper plates of the heated imaging platform. A small quantity of Aquasonic gel was applied to the paws before taping to ensure a full ECG trace. Both long and short-axis heart views were acquired in B and M mode and analyzed by the "LV analysis" software tool (VisualSonics). Each mouse was kept warm using a heated platform and warm Aquasonic ultrasound gel while scanning. The gel was placed on the mid-abdomen to the upper ventral side of the mice. Clips were taken of the long and short axis of each mouse's hearts in both B and M mode. After the scanning was completed, Vevo Lab Software analysis was used to analyze the images and perform the Cardiac Package of LV Trace on the long and short axis.

**Statistical analysis:**
Statistical tests were performed and analyzed using Microsoft Excel and ggplot2. P-value was calculated by two-tailed paired Student's t-test. Significance was defined as $P < 0.05$. All data are presented as mean ± SEM.

**References:**

Anders S, Pyl PT, Huber W (2015). HTSeq--a Python framework to work with high-throughput sequencing data. Bioinformatics 15, 166-9.

Conrad T and Ørom AU (2017). Cellular Fractionation and Isolation of Chromatin-Associated RNA. Methods Mol Biol 1468, 1-9.

Dobin A, Davis CA, Schlesinger F, Drenkow J, Zaleski C, Jha S, Batut P, Chaisson M, Gingeras TR (2013). STAR: ultrafast universal RNA-seq aligner. Bioinformatics 29, 15-21.

Han Y, Slivano OJ, Christie CK, Cheng AW, Miano JM (2015). CRISPR-Cas9 genome editing of a single regulatory element nearly abolishes target gene expression in mice--brief report. Arterioscler Thromb Vasc Biol 35, 312-5.

Janky, R., Verfaillie, A., Imrichova, H., Van de Sande, B., Standaert, L., Christiaens, V., Hulselmans, G., Herten, K., Naval Sanchez, M., Potier, D., et al. (2014). iRegulon: from a gene list to a gene regulatory network using large motif and track collections. PLoS Comput Biol 10, e1003731.

Klattenhoff, C.A., Scheuermann, J.C., Surface, L.E., Bradley, R.K., Fields, P.A., Steinhauser, M.L., Ding, H., Butty, V.L., Torrey, L., Haas, S., et al. (2013). Braveheart, a long noncoding RNA required for cardiovascular lineage commitment. Cell 152, 570-583.

Love MI, Huber W, Anders S (2014). Moderated estimation of fold change and dispersion for RNA-seq data with DESeq2. Genome Biol 15, 550.

Luo W and Brouwer C (2013). Pathview: an R/Bioconductor package for pathway-based data integration and visualization. Bioinformatics 29, 1830-1.

Luo W, Friedman MS, Shedden K, Hankenson KD, Woolf PJ (2009). GAGE: generally applicable gene set enrichment for pathway analysis. BMC Bioinformatics 10, 161.

Figure S1

A

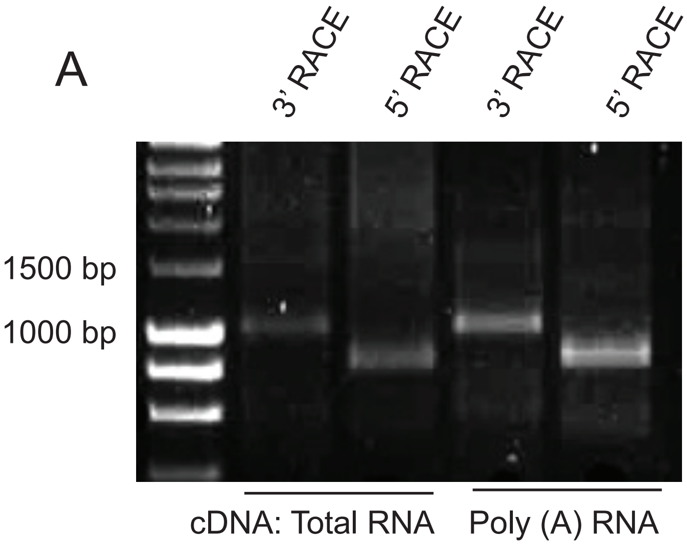


B

| Method | Candidates | Values | Protein-coding cutoff |
|---|---|---|---|
| CPAT | *Platr4* | 0.02312 | >0.44 |
| CPC | *Platr4* | -1.09113 | >0 |

C

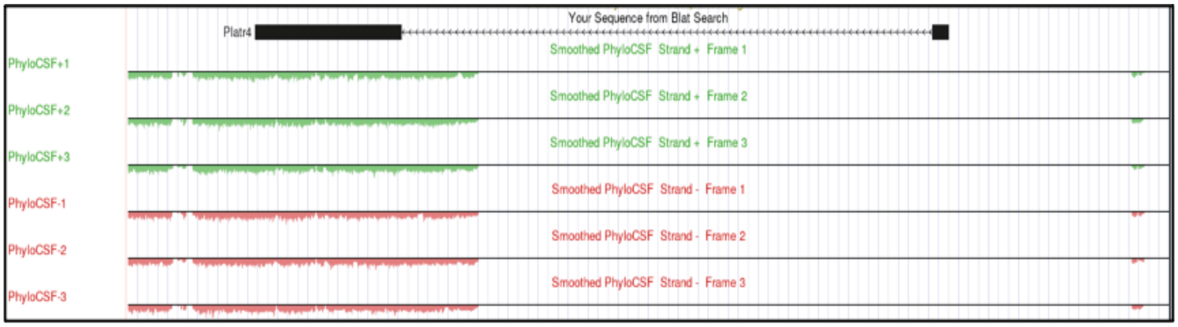


D

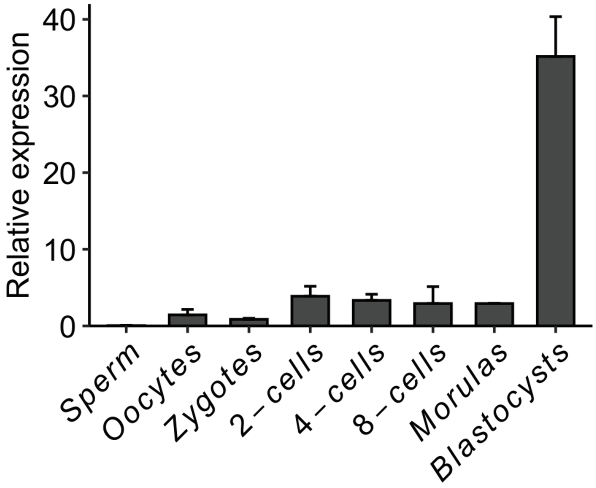


E

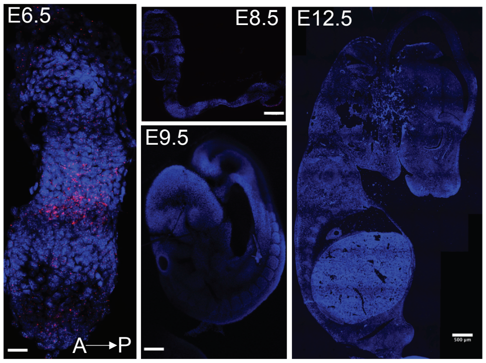

### Figure S2

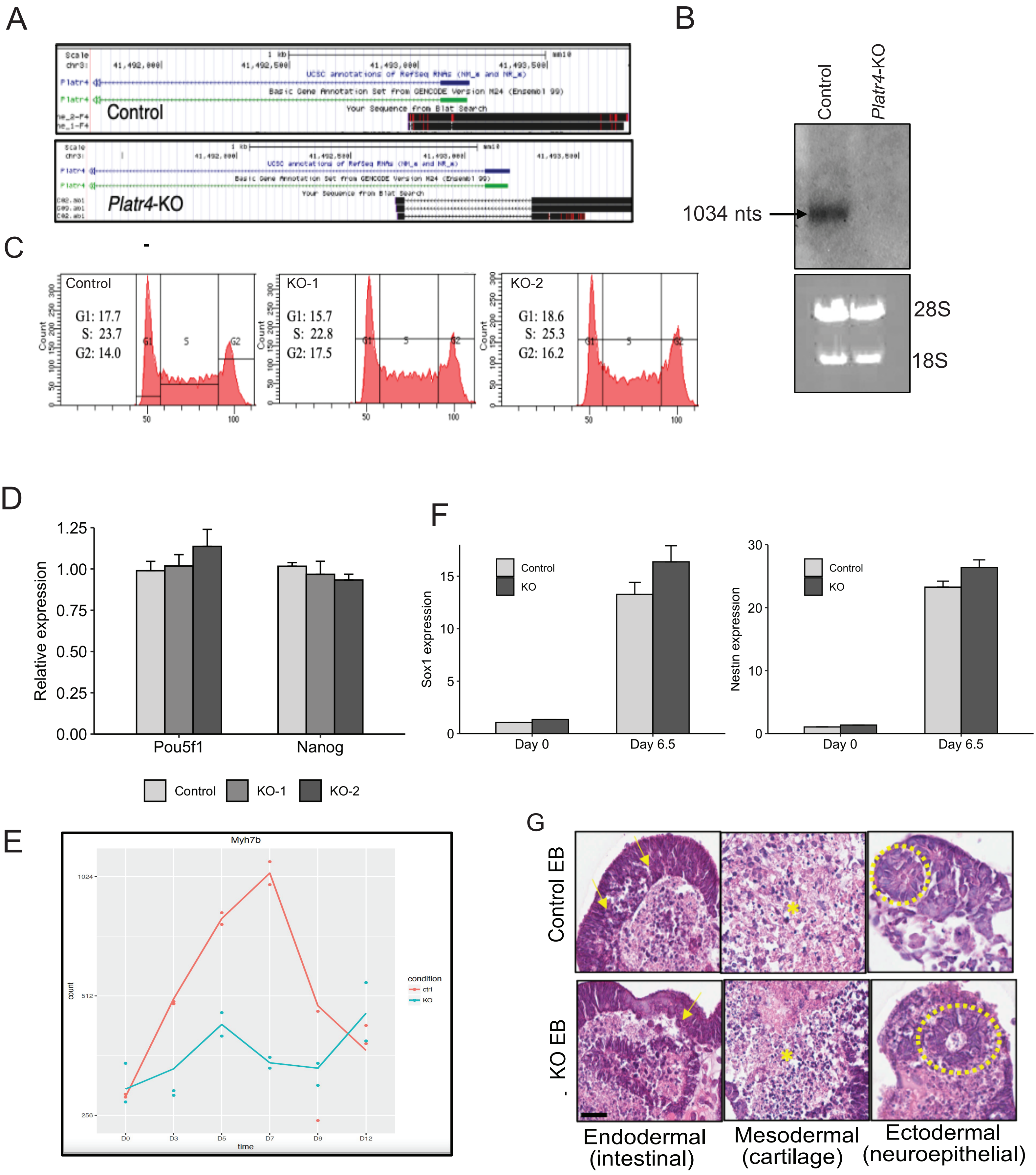

A
Control
Platr4-KO
B
Control
Platr4-KO
1034 nts
28S
18S
C
Control
G1: 17.7
S: 23.7
G2: 14.0
KO-1
G1: 15.7
S: 22.8
G2: 17.5
KO-2
G1: 18.6
S: 25.3
G2: 16.2
Count
D
Relative expression
Pou5f1
Nanog
Control
KO-1
KO-2
F
Sox1 expression
Nestin expression
Control
KO
Day 0
Day 6.5
E
Myh7b
count
time
condition
ctrl
KO
G
Control EB
KO EB
Endodermal (intestinal)
Mesodermal (cartilage)
Ectodermal (neuroepithelial)

### Figure S3

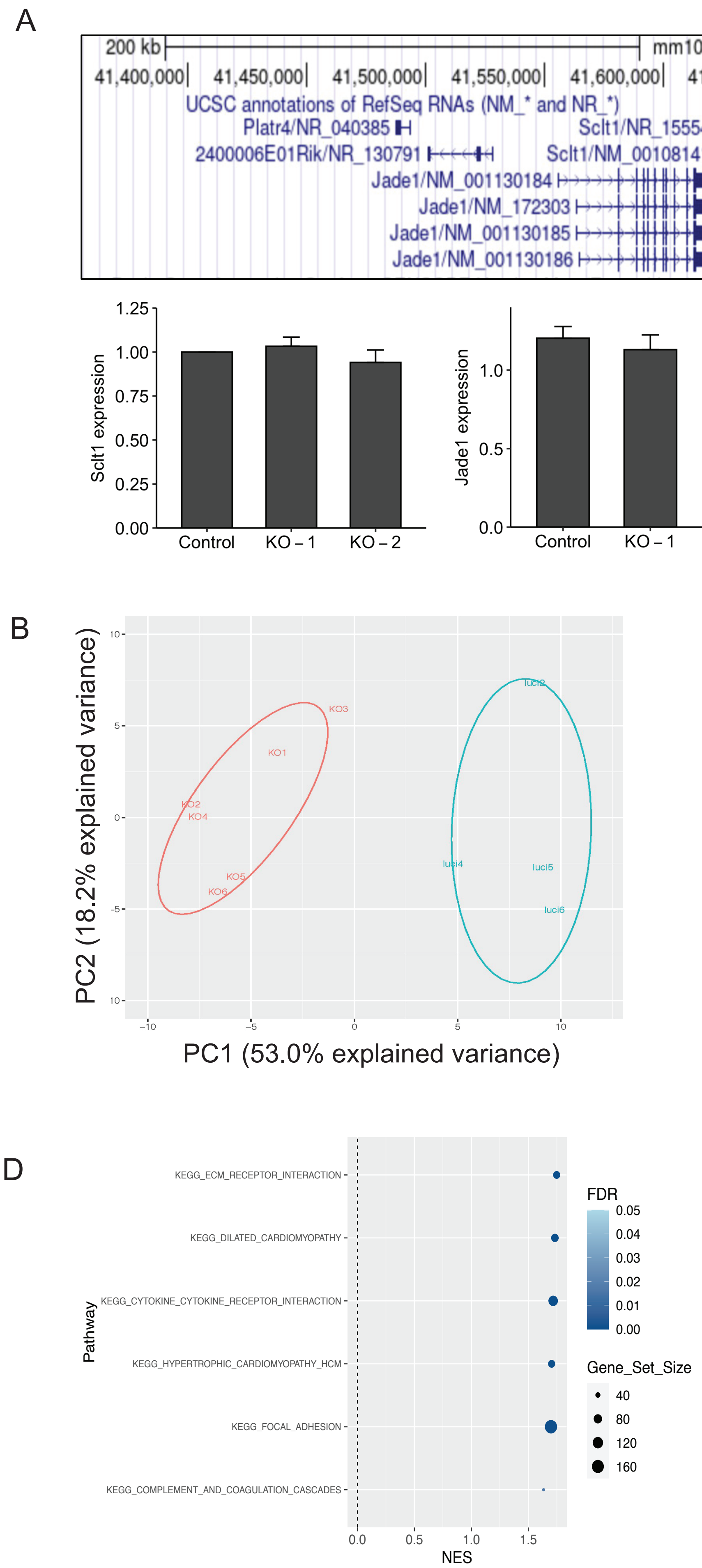

A
200 kb
mm10
41,400,000
41,450,000
41,500,000
41,550,000
41,600,000
41,650,000
41,700,000
41,750,000
41,800,000
UCSC annotations of RefSeq RNAs (NM_* and NR_*)
Platr4/NR_040385
2400006E01Rik/NR_130791
Jade1/NM_001130184
Jade1/NM_172303
Jade1/NM_001130185
Jade1/NM_001130186
Sclt1/NR_155540
Sclt1/NM_001081411
D3Ertd751e/NM_001291048
D3Ertd751e/NM_001099785
D3Ertd751e/NM_027271
D3Ertd751e/NM_028667
Sclt1 expression
1.25
1.00
0.75
0.50
0.25
0.00
Control
KO – 1
KO – 2
Jade1 expression
1.0
0.5
0.0
D3Ertd751 expression
0.4
0.3
0.2
0.1
0.0
B
PC2 (18.2% explained variance)
PC1 (53.0% explained variance)
KO1
KO2
KO3
KO4
KO5
KO6
luci2
luci4
luci5
luci6
10
5
0
-5
-10
–10
–5
0
5
10
C
Enrichment score
NES = 2.11
FDR = 0.001
Pluripotent genes
D
Pathway
KEGG_ECM_RECEPTOR_INTERACTION
KEGG_DILATED_CARDIOMYOPATHY
KEGG_CYTOKINE_CYTOKINE_RECEPTOR_INTERACTION
KEGG_HYPERTROPHIC_CARDIOMYOPATHY_HCM
KEGG_FOCAL_ADHESION
KEGG_COMPLEMENT_AND_COAGULATION_CASCADES
0.0
0.5
1.0
1.5
NES
FDR
0.05
0.04
0.03
0.02
0.01
0.00
Gene_Set_Size
40
80
120
160

Figure S4

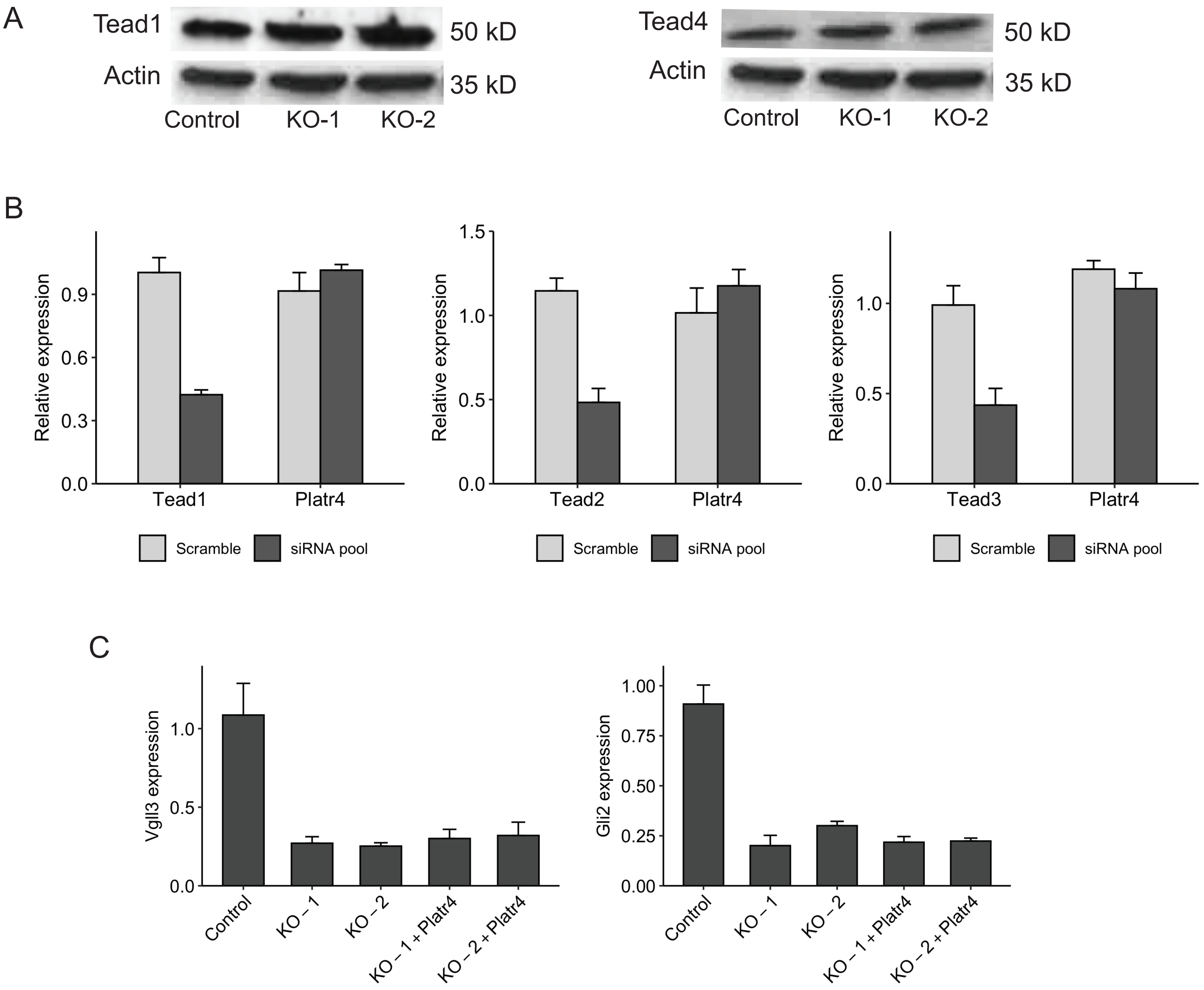
A
Tead1
50 kD
Actin
35 kD
Control
KO-1
KO-2
Tead4
50 kD
Actin
35 kD
Control
KO-1
KO-2
B
Relative expression
Tead1
Platr4
Tead2
Platr4
Tead3
Platr4
Scramble
siRNA pool
C
Vgll3 expression
Gli2 expression
Control
KO – 1
KO – 2
KO – 1 + Platr4
KO – 2 + Platr4

### Figure S5

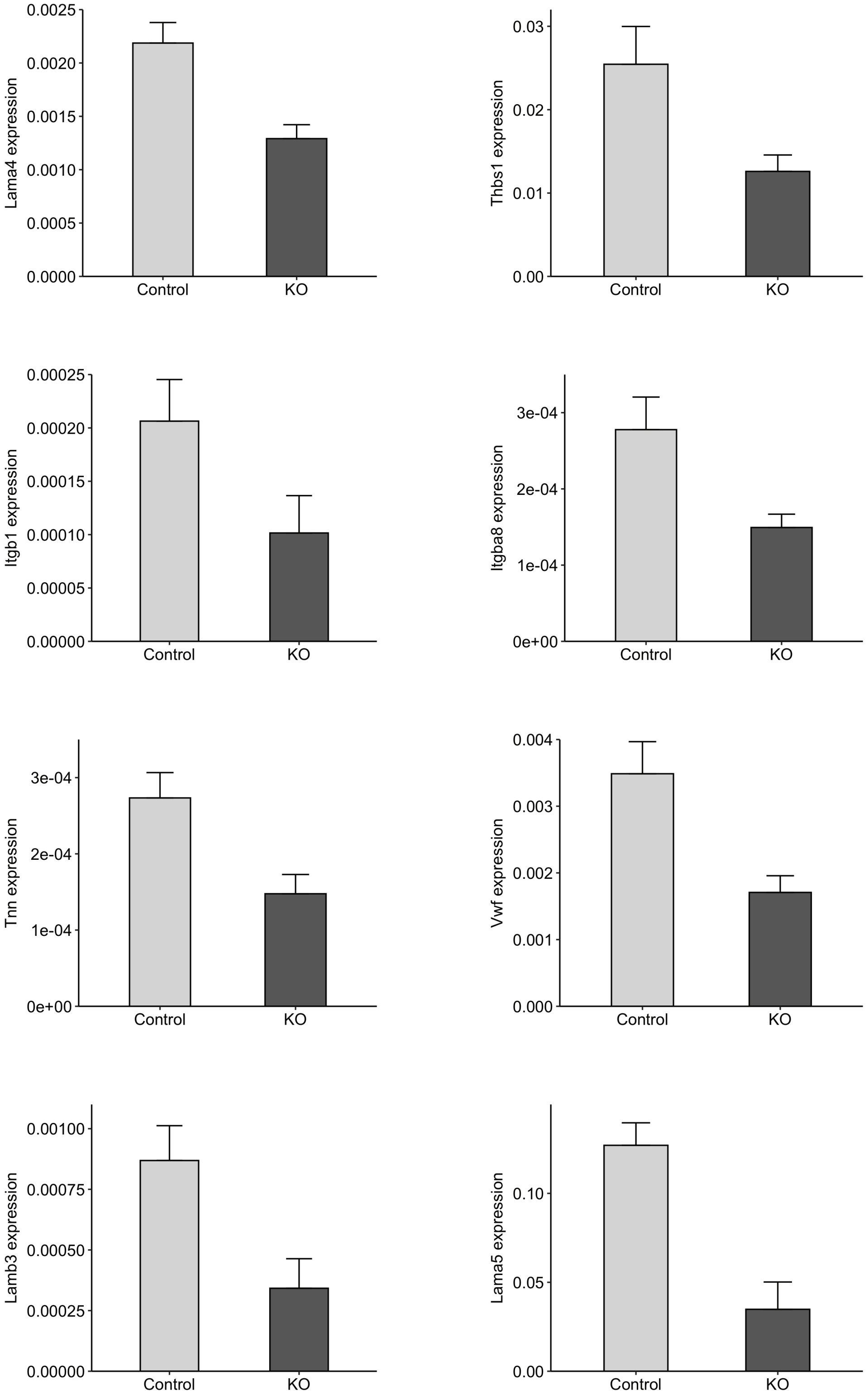
Lama4 expression
0.0025
0.0020
0.0015
0.0010
0.0005
0.0000
Control
KO
Thbs1 expression
0.03
0.02
0.01
0.00
Control
KO
Itgb1 expression
0.00025
0.00020
0.00015
0.00010
0.00005
0.00000
Control
KO
Itgba8 expression
3e-04
2e-04
1e-04
0e+00
Control
KO
Tnn expression
3e-04
2e-04
1e-04
0e+00
Control
KO
Vwf expression
0.004
0.003
0.002
0.001
0.000
Control
KO
Lamb3 expression
0.00100
0.00075
0.00050
0.00025
0.00000
Control
KO
Lama5 expression
0.10
0.05
0.00
Control
KO

Figure S6

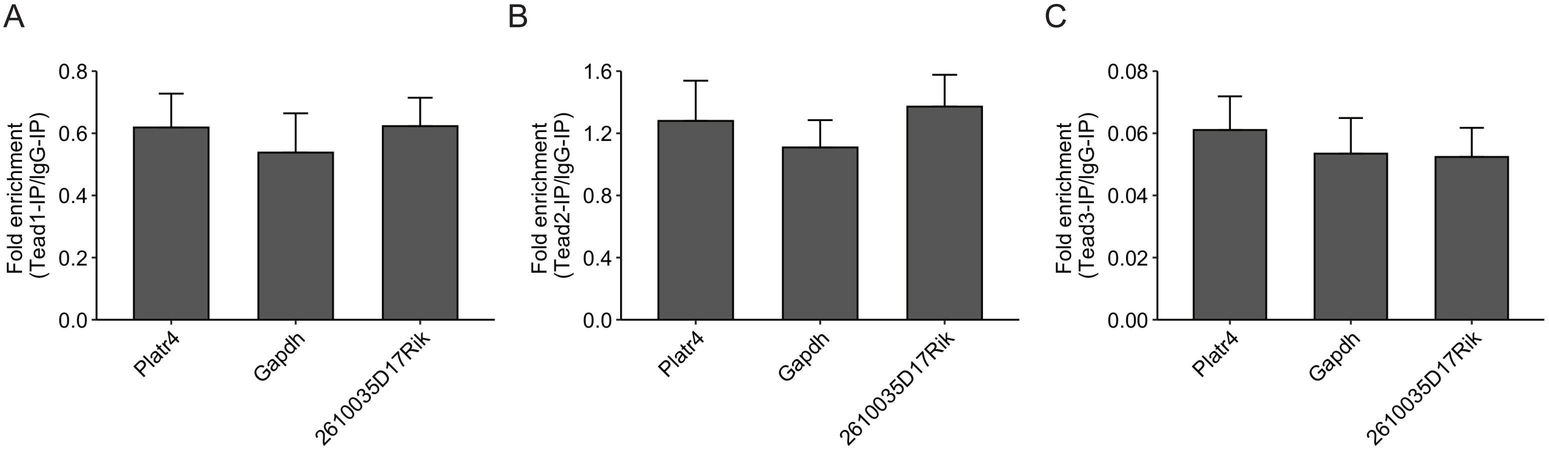
A
B
C
Fold enrichment
(Tead1-IP/IgG-IP)
Fold enrichment
(Tead2-IP/IgG-IP)
Fold enrichment
(Tead3-IP/IgG-IP)
0.8
0.6
0.4
0.2
0.0
1.6
1.2
0.8
0.4
0.0
0.08
0.06
0.04
0.02
0.00
Platr4
Gapdh
2610035D17Rik

### Figure S7

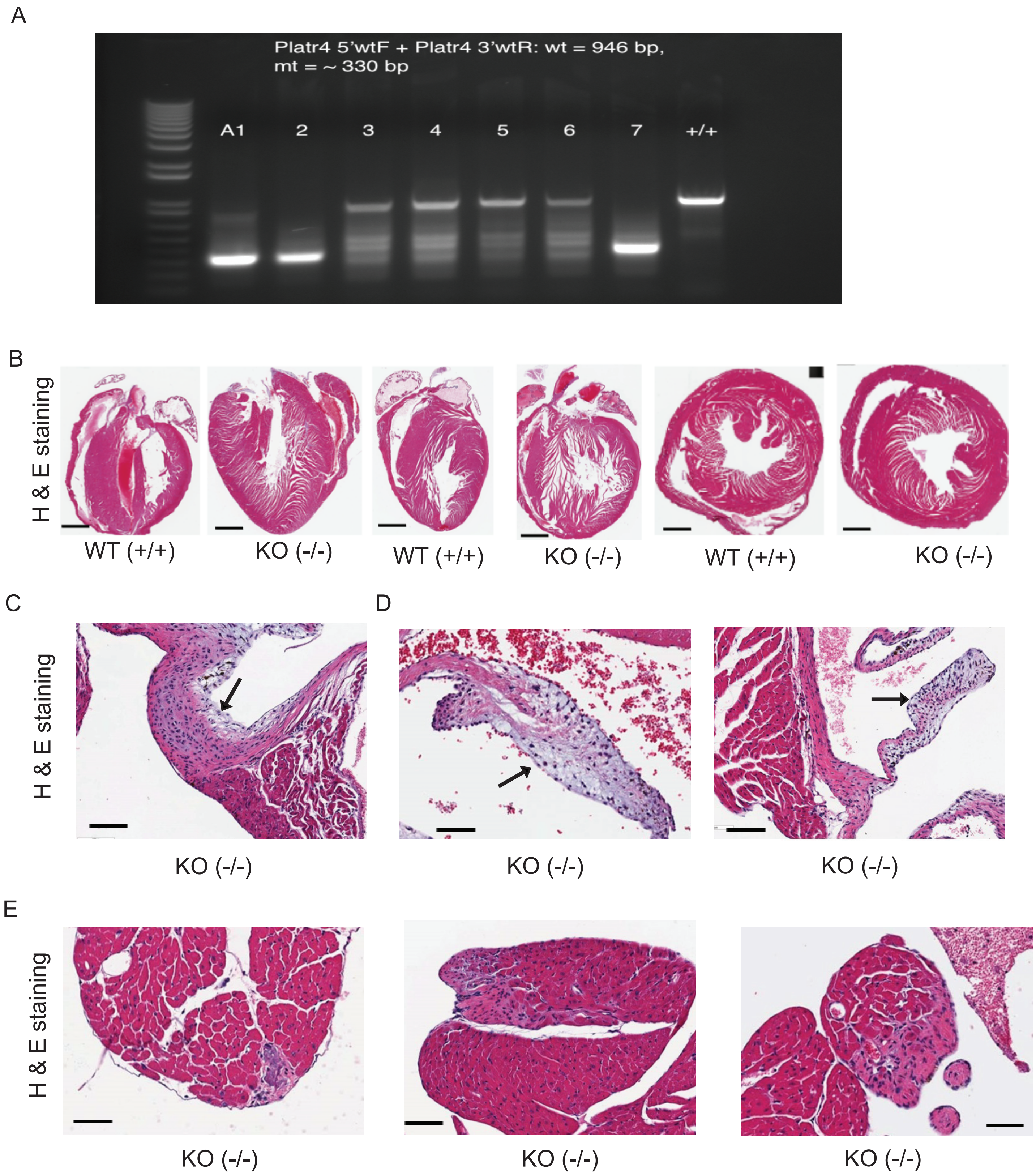
A
Platr4 5'wtF + Platr4 3'wtR: wt = 946 bp,
mt = ~ 330 bp
A1
2
3
4
5
6
7
+/+
B
H & E staining
WT (+/+)
KO (-/-)
WT (+/+)
KO (-/-)
WT (+/+)
KO (-/-)
C
D
H & E staining
KO (-/-)
KO (-/-)
KO (-/-)
E
H & E staining
KO (-/-)
KO (-/-)
KO (-/-)

**Supplemental Figure Legend**

### *Platr4* lncRNA regulates mesoderm/endoderm differentiation through the TEAD4-CTGF axis.

Rasmani Hazra, Lily Brine, Libia Garcia, Brian Benz, Napon Chirathivat, Michael Shen, John Erby Wilkinson, Scott K. Lyons, David L. Spector

***Figure S1: Platr4 is an ESC-specific nuclear enriched lncRNA***

(A) 5′ and 3′ rapid amplification of cDNA ends (RACE) were performed to identify the full-length *Platr4* transcript. Three independent experiments were performed, and a representative gel image is shown. (B) Protein coding potential analysis of *Platr4* RNA transcript using Coding-Potential Assessment Tool (CPAT) and Coding Potential Calculator (CPC). (C) Screenshot of Platr4 genomic locus from UCSC genome browser shows PhyloCSF tracks with no coding potential. (D) Relative expression of Platr4 in sperm, oocytes, and pre-implanted embryos measured by qRT-PCR (n=3, independent experiments). (E) Single-molecule RNA-FISH (red dots) in post-implanted embryos (E6.5 to 12.5). A and P stands for anterior and posterior. Scale bars are 500 μm.

***Figure S2: Platr4 is essential for ESC differentiation***

(A) A snapshot of BLAT results showing the CRISPR/Cas9-mediated genome editing *Platr4*-knockout (KO) vs. control clones over the promoter region of the Platr4 gene locus by aligning Sanger sequencing results on the UCSC genome browser. (B) Deletion of Platr4 RNA in ESCs was measured by Northern blot using total RNA. 28S and 18S bands are the indicators of the quality of the RNA. (C) Cell cycle analysis of control and Platr4-KO clones were assessed using flow cytometry to compare the percentage of cells in each cell cycle stage (n=2 independent experiments). (D) Relative expression of pluripotent master transcription factors (*Pou4f2* and *Nanog*) in control vs. Platr4-KO ESCs, measured by qRT-PCR. (E) The relative level of neuroectodermal lineage markers (*Sox1*, *Nestin*) was measured by qRT-PCR using control vs. *Platr4*-KO EBs at different time points upon differentiation. (F) Expression of Myosin heavy chain 7b (Myh7b) in control vs. Platr4-KO EBs at different time points upon differentiation. (G)

Hematoxylin and eosin (H&E)-stained EB sections of control vs. Platr4-KO clones showed three germ layers. Scale bar, 200 µm. All experiments were performed in triplicate. *$p < 0.05$ (student's t-test).

***Figure S3: Platr4 regulates mesoderm/endoderm lineage specification***

(A) qRT-PCR analysis of neighboring protein coding genes (*Sclt1*, *Jade1*, and *D3Ertd751*) within 100kb span of Platr4 locus in control vs. *Platr4*-KO ESCs. (B) Principal component analysis (PCA) plot of control and Platr4-KO clones in RNA-seq data. (C) Gene set enrichment analysis profiles of DE genes to measure pluripotency. The gene lists have shown in supplemental table 3. (D) Analysis of significant GO terms in downregulated DEG in *Platr4*-KO EBs.

***Figure S4: Platr4 functions upstream of connective tissue growth factor (Ctgf)***

(A) Western blot analyses of Tead1 and Tead4 in control and *Platr4*-KO ESCs. Actin was used as an internal control. (B) qRT-PCR analysis of *Platr4* level in ESCs using Tead1, Tead2, and Tead3-siRNA and control siRNA. (C) Tead4 downstream target genes (*Vgll3* and *Gli2*) was not rescued upon ectopic expression of *Platr4* in *Platr4*-KO cells as determined by qRT-PCR. All experiments were performed in triplicate. Results are mean ± SD (n=3) *$p < 0.05$ (student's t-test).

***Figure S5: Directed cardiac differentiation***

Significant downregulated of ECM-genes in *Platr4*-KO compared to control ESCs as measured by qRT-PCR. Values are mean ± SD (n=3) *$p < 0.05$ (two-tailed student's t-test).

***Figure S6: Platr4 interacts with Tead4***

(A, B & C) RIP assay confirmed that *Platr4* did not interact with Tead1, Tead2, and Tead3 using respective antibodies. Fold enrichment of *Patr4* over IgG signal was shown. *Gapdh*, and *2610035D17Rik* (lncRNA) transcripts were used for controls. Values are mean ± SD (n=3) *p < 0.05 (two-tailed student's t-test).

***Figure S7: Phenotype of Platr4-KO mice***

(A) PCR genotyping of three founder pups (A1, A2, and A7) was shown here. (B) H&E staining (both longitudinal and horizontal sections) for the adult heart of WT and Platr4-KO mice. (C) H&E staining adult Platr4-KO heart was showing valve defect (fibro cartilaginous metaplasia). Scale bar: 200 µm. (D) H&E staining adult Platr4-KO heart was showing mucinous valve degeneration. Scale bar: 100 µm. (E) H&E staining adult Platr4-KO heart was showing myocardial atrophy and fibrosis. Scale bar: 200 µm.
